# Dynamical Modulation of Hippocampal Replay Sequences through Firing Rate Adaptation

**DOI:** 10.1101/2024.09.13.612895

**Authors:** Zilong Ji, Tianhao Chu, Xingsi Dong, Changmin Yu, Daniel Bush, Neil Burgess, Si Wu

## Abstract

During periods of immobility and sleep, the hippocampus generates diverse self-sustaining sequences of “replay” activity, exhibiting stationary, diffusive, and super-diffusive dynamical patterns. However, the neural mechanisms underlying this diversity in hippocampal sequential dynamics remain largely unknown. Here, we propose such a mechanism demonstrating that modulation of firing rate adaptation in a continuous attractor model of place cells causes the emergence of different types of replay. Our model makes several key predictions. First, more diffusive replay sequences positively correlate with longer theta sequences across animals (both reflecting stronger adaptation). Second, replay diffusivity varies within an animal across behavioural states that affect adaptation (such as wake and sleep). Third, increases in neural excitability, incorporated with firing rate adaptation, reduce the step size of decoded movements within individual replay sequences. We provide new experimental evidence for all three predictions. These insights suggested that the diverse replay dynamics observed in the hippocampus can be reconciled through a simple yet effective neural mechanism, shedding light on its role in hippocampal-dependent cognitive functions and its relationship to other aspects of hippocampal electrophysiology.

## 1 Introduction

The hippocampus plays a pivotal role in various cognitive processes, including navigational learning [Keefe and Nadel, 1978, Morris et al., 1982], goal-directed decision making [Johnson and Redish, 2007, Wikenheiser and Redish, 2015, Kay et al., 2020], and episodic memory [Scoville and Milner, 1957, Olton and Samuelson, 1976, Steele and Morris, 1999]. These cognitive processes require the temporal coding of relationships between events and/or locations, with hippocampal sequential activity hypothesized to support these computations [Wilson and McNaughton, 1994, Skaggs et al., 1996, Foster and Wilson, 2007, Diba and Buzsáki, 2007, Karlsson and Frank, 2009, Gupta et al., 2010]. Empirical studies have identified two distinct yet interrelated types of hippocampal sequences. First, during active exploration, place cells fire in sequences within individual LFP theta cycles, termed theta sequences (Fig. 1a&b) [Skaggs et al., 1996, Foster and Wilson, 2007]. Nested within longer behavioral timescales, theta sequences compress spiking intervals between successive place cells to within dozens of milliseconds, thereby facilitating Hebbian synaptic plasticity [O’Keefe and Recce, 1993, Jensen and Lisman, 1996, Magee and Johnston, 1997], which is essential for the initial formation of memory traces [Drieu et al., 2018, Muessig et al., 2019, Drieu and Zugaro, 2019]. Furthermore, their forward-directed nature positions them as suitable for spatial planning and decision-making [Johnson and Redish, 2007, Wikenheiser and Redish, 2015, Kay et al., 2020]. Second, during periods of immobility and sleep, behavioral sequences encoded in the hippocampus can be reactivated during “replay” events (Fig. 1c&d). Similar to theta sequences, replay sequences also occur as compressed sequences lasting tens to hundreds of milliseconds, associated with sharp-wave ripple events (SWRs) in the local field potential (LFP) [Lee and Wilson, 2002]. Replay sequences vary in form, for instance, forward/reverse replay of the current experience [Diba and Buzsáki, 2007, Foster and Wilson, 2006], remote replay of past experiences [Karlsson and Frank, 2009], and preplay of upcoming events [Dragoi and Tonegawa, 2011]. These variations have been taken as evidence of distinct mechanisms and cognitive functions, such as memory consolidation and goal-directed planning.

**Figure 1:**
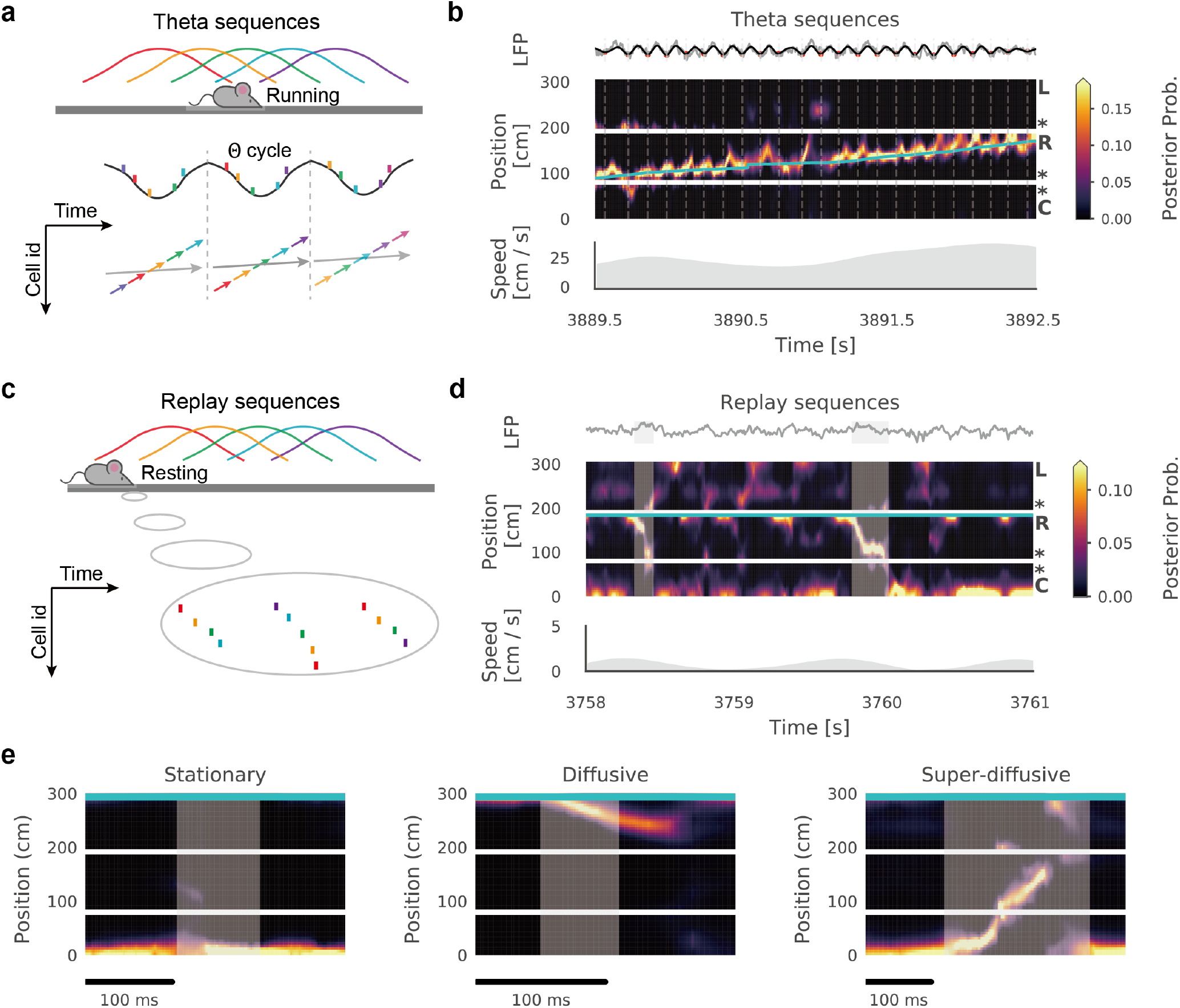
Sequential dynamics in the hippocampus. **(a)**, schematic of theta phase precession at the single neuron level and theta sequences at the population level. **(b)**, example theta sequences in the experimental data when an animal performed a W-track spatial alternation task (see Fig.S1 for more details). Top panel: theta-band (5-11 Hz) filtered LFP. Middle panel: decoded position; linearized from the central arm (C) to the right arm (R) and then to the left arm (L). The posterior probability is plotted as color values with the blue line representing the actual location of the animal. Bottom panel: running speed of the animal. **(c)**, schematic of replay sequences. **(d)**, example replay sequences in the experimental data when the animal stopped at the end of the right arm (marked by the blue line). Grey areas indicates periods of detected SWRs. **(e)**, Diverse replay dynamics. From left to right: stationary (decoded positions stay at a single location), diffusive (decoded positions slowly propagate), and superdiffusive (decoded positions jump to another location) replay sequences. Grey areas indicate periods of detected SWRs.

While theta sequences regularly sweep through the animal’s location during active running, many replay sequences do not exhibit steady propagation and instead display a wide variety of dynamics (Fig. 1e) [Denovellis et al., 2021]. For example, certain replay sequences consistently represent a single location away from the animal, potentially indicating reward wells or choice points [Yu et al., 2017]. During task-engaged periods in spatial memory tasks, replay sequences exhibit “jumping” movements [Pfeiffer and Foster, 2015], alternating between single and spatially discontinuous locations, a phenomenon termed super-diffusive dynamics. Conversely, during sleep periods following a random foraging task, replay sequences manifest Brownian diffusion dynamics [Stella et al., 2019], which do not directly mirror behavioral trajectories. Besides these task differences, studies have also shown that replay dynamics undergo developmental changes: representing single locations in pre-weaning stages and gradually evolving into trajectory-like sequences after weaning [Muessig et al., 2019, Farooq and Dragoi, 2019]. These findings illustrate that multiple factors influence the detailed dynamics of replay sequences, raising important questions about their underlying mechanisms. Specifically, what conditions are necessary and sufficient for the generation of replay sequences in the hippocampus? What factors dictate the emergence of specific replay dynamics, and how does the brain adjust and adapt these mechanisms to different circumstances? Additionally, how are different hippocampal sequences, particularly theta sequences during active behavior and replay sequences during rest, interconnected?

To elucidate the neural mechanisms responsible for generating diverse replay dynamics in the hippocampus, we develop a theoretical framework conceptualizing the hippocampal place cell assembly as a continuous attractor network (CAN). In this model, spatial representation in the hippocampus is embodied as an activity bump within the CAN, with replay sequences emerging as spontaneous movements of this bump. Here, we demonstrate, both theoretically and empirically, that firing rate adaptation serves as an intrinsic mechanism that destabilizes the activity bump, with changes in the strength of adaptation leading to a variety of replay-like dynamics, including the stationary, Brownian-diffusive, and super-diffusive sequences observed in empirical studies. Beyond providing theoretical explanations for replay diffusivity, we expanded our research by making predictions and validating them with experimental data.

Initially, inspired by studies demonstrating the concurrent development of theta and replay sequences [Muessig et al., 2019, Drieu et al., 2018], we explore how replay dynamics are associated with theta sequences during active behaviors, as theta sequences can also be captured by this model [Chu et al., 2024]. Our model predicts that more diffusive replay is correlated with longer theta sequences across animals (both reflecting stronger adaptation), a relationship subsequently confirmed by experimental data. Next, inspired by research showing different levels of neurotransmitters such as acetylcholine, dopamine, and noradrenaline across different brain states [Samanta et al., 2020], which potentially affect firing rate adaptation [Yoshida et al., 2013, Barkai and Hasselmo, 1994, Madison and Nicoll, 1984], we examine how replay dynamics vary within the same animal during different brain states. Our model predicts that varying adaptation strength in the simulated hippocampal network leads to different replay dynamics. Inspired by this, we find that within individual animals, awake replay sequences are more diffusive than subsequent sleep replay sequences. Finally, inspired by a study demonstrating that replay trajectories display interleaved local- and long-jump movements [Pfeiffer and Foster, 2015], we investigate the detail structures of individual replay sequences. Our model predicts that increases in neural excitability, in cooperation with firing rate adaptation, reduce the step size of replay trajectories. This reveals a detailed dynamical structure within individual replay sequences, which was further confirmed by empirical data. Overall, our study proposes a fundamental mechanism integrating diverse hippocampal sequences observed empirically, making predictions at multiple hierarchical levels and validating them with experimental data. Consequently, these findings advance our understanding of sequence generation in the hippocampus and elucidate the computational principles underpinning its functions.

## 2 Results

### 2.1 A continuous attractor network with firing rate adaptation for the hippocampal place cell assembly

To understand the underlying mechanism of replay dynamics in the hippocampus, we implemented a conventional continuous attractor network (CAN) model [McNaughton et al., 2006] of the hippocampal place cell population (see Methods. 4.3 for mathematical details). In particular, neurons in the CAN model are arranged based on the locations of their firing fields in the environment (Fig. 2a), with the dynamics expressed as

**Figure 2:**
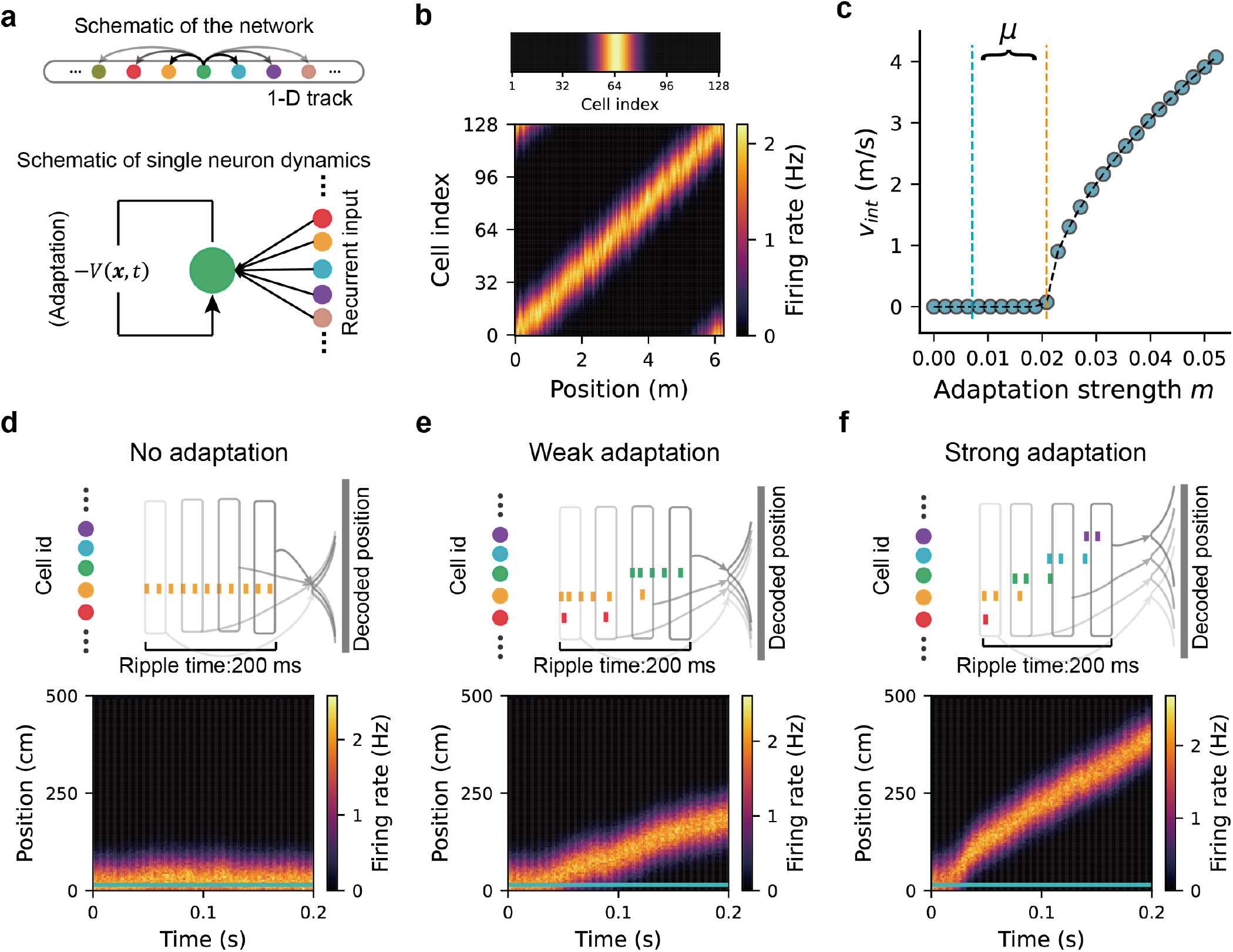
Hippocampal place cells modelled as a continuous attractor network (CAN) with firing rate adaptation. **(a)**, top panel: schematic of the CAN for modeling the place cell network in the linear track environment. Neurons, indicated by dots with different colors, are rearranged according to the location of their firing field in the environment. Bottom panel: single neuron dynamics in the CAN with recurrent input from other neurons and firing rate adaptation illustrated as a negative feedback to the neuron. **(b)**, top panel: bump-like network activity in the simulated CAN. Bottom panel: localized firing field for all the neurons in the CAN. **(c)**, the intrinsic travelling speed of the activity bump as a function of the adaptation strength. The orange line marks the transition boundary below which the activity bump is stationary (see Methods. 4.5 for theoretical derivation). The distance of the adaptation strength (exampled as the blue line) to the transition boundary is marked as a parameter of *µ* in theoretical analysis below (Fig. 3a&b and Eq. (4)). **(d)**, replay-like sequences with no adaptation. Top panel: schematic of generating stationary sequences under no adaptation. Each vertical bar represents a spike event with the color matching the identity of the place cell. Each box represents a time bin with decoded position shown on the right side. Bottom panel: population activity in the CAN. The animal stays at the bottom of the linear track environment, indicated by the horizontal blue line. **(e)**, replay-like sequences with weak adaptation. **(f)**, replay-like sequences with strong adaptation.

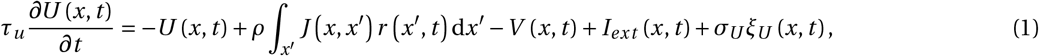

where *U* (*x, t*) denotes the synaptic input to the cell at location *x*, and *r* (*x, t*) is the corresponding firing rate. Each place cell forms recurrent connections with all other cells through excitatory synapses *J (x, x*^′)^ (scaled by *ρ*), where the connection strength diminishes proportionally to the distance between the place fields of two cells. This translation invariant recurrent connection can be formed by a Gaussian random walk in the environment [Stachenfeld et al., 2017, Yu et al., 2020]. Additionally, the network experiences global feedback inhibition which limits the total neural activity (Eq. (7) in Methods. 4.3). These network configurations lead to the emergence of an activity bump as the stable state of the network, accompanied by localized firing fields of individual cells (see Fig. 2b and Methods. 4.4 for the theoretical deriva-tion). The CAN model also receives a location dependent sensory input *I*_*ext*_ (*x, t*), which is active during animals’ running (see Results. 2.4.1 below) but absent during immobility. Furthermore, the model experiences internally generated noise *σ*_*U*_ *ξ*_*U*_ (*x, t*). In the absence of input other than network noise, the bump location remains either stable or undergoes a random drift in position, resembling the experimental finding [Stella et al., 2019].

Importantly, neurons in the network exhibit firing rate adaptation (see Fig. 2a), a general feature of neural responses in the brain [Hopfield, 2010, Itskov et al., 2011, Fuhrmann et al., 2002, Benda and Herz, 2003]. While this property can be attributed to various biophysical mechanisms, the common feature involves a form of slow negative feedback *V* (*x, t*) to the cell’s excitability, which is expressed as:

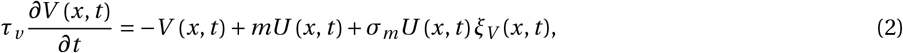

where *τ*_*v*_ is the adaptation time constant much larger than *τ*_*u*_ . *m* is the adaptation strength and *ξ*_*V*_ (*x, t*) is the noise with *σ*_*m*_ the noise level in firing rate adaptation. This destabilization makes the activity bump move with an intrinsic speed increasing with the adaptation strength (see Fig. 2c and Methods. 4.5 for the theoretical derivation). Specifically: 1) when there is no adaptation (or adaptation strength below the threshold showed in Fig. 2c), the firing frequencies of active cells remain unchanged, and the activity bump remains stationary (Fig. 2d); 2) when the adaptation strength is weak (but above the threshold in Fig. 2c), the firing frequencies of active cells gradually decrease, leading to increased firing of nearby cells due to the strong recurrent connection with the preceding active cells and the competition mediated by global inhibition. Consequently, the activity bump moves slowly to the location of nearby cells (Fig. 2d); and 3) when the adaptation strength is strong (well above the threshold), the firing frequencies of active cells decrease rapidly, causing nearby neurons to quickly increase their firing. This rapid succession of firing events results in rapid movement of the activity bump (Fig. 2e).

### 2.2 Firing rate adaptation strength accounts for the diverse diffusivity of replay sequences in the CAN model

We have shown that firing rate adaptation leads to the spontaneous propagation of activity at a constant speed, which explains a portion of the sequential dynamics observed during sharp wave-ripple complexes (SWRs) [Denovellis et al., 2021]. In this context, we hypothesize that the interplay between network noise and the strength of adaptation could account for the diverse replay dynamics observed in the hippocampus. To test this hypothesis, we conducted theoretical analyses using two distinct approaches. The first approach aims to mechanistically understand the sequential dynamics in the CAN by solving the dynamical system. The second approach seeks to comprehend the sequential dynamics normatively by performing spectral analysis on the transition matrix within the CAN. These complementary analytical approaches provide profound insights into the genesis of heterogeneous replay profiles.

#### 2.2.1 Dynamical system analysis from a mechanistic viewpoint

The first analysis involves treating the CAN model as a dynamical system and deriving the condition for generating diverse replay dynamics based on the adaptation strength and noise level. We begin by providing two intuitive examples to illustrate this mechanism. For the generation of Brownian-diffusive sequences, we configure the model with weak adaptation and significant network noise. In this scenario, the activity bump undergoes random drift due to the noise rather than directional movement due to the adaptation. Conversely, to generate super-diffusive sequences, we configure the model with strong adaptation (choosing a value near the travelling boundary in Fig. 2c) and medium network noise. In this case, random noise causes the adaptation strength to fluctuate around the travelling boundary. Consequently, the bump location undergoes intermittent switches between local movement (when the adaptation strength stays below the boundary) and long-jump movement (when the adaptation strength surpasses the boundary), resembling the features of super-diffusive dynamics.

To fully characterize the diffusivity of the sequential dynamics arising from the interplay between network noise and adaptation strength, we quantified bump trajectories as a power law distribution of step size, which is:

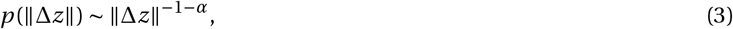

where Δ*z* is the step size of the activity bump in Δ*t* with Δ*t* → 0, and *α* represents the power-law exponent. When *α* ≥ 2, the bump movement trajectory is characterized as Brownian motion. When 0 < *α* < 2, the bump movement trajectory is characterized as super-diffusion. Note that we ignore *α* ≤ 0 since this corresponds to both infinite values of the mean and variance of the step size in theory. Via theoretical analysis (Methods. 4.6 & 4.7), we found that the power-law exponent depends on two key parameters in our model, i.e., the adaptation strength and the noise amplitude, with the form as:

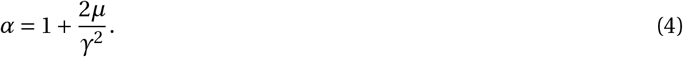

Here *µ* measures the difference between the adaptation strength and the transition boundary shown in Fig. 2c, and *γ* measures the normalized noise level (related to *σ*_*m*_ in Eq. 2, see Methods. 4.6 for more details). The relationship between the power-law exponent and the two variables is visualized in a phase diagram (Fig. 3a), which shows that 1) small *µ* (adaptation strength is close to the transition boundary) and large *γ* (high network noise level) lead to super-diffusive dynamics; 2) large *µ* (adaptation strength is far below the transition boundary) and medium *γ* (medium network noise level) lead to Brownian-diffusive dynamics; 3) large *µ* (adaptation strength is far below the transition boundary) and small *γ* (small network noise level) lead to stationary dynamics. These analytical results were further verified by numerical simulations (Fig. 3b).

**Figure 3:**
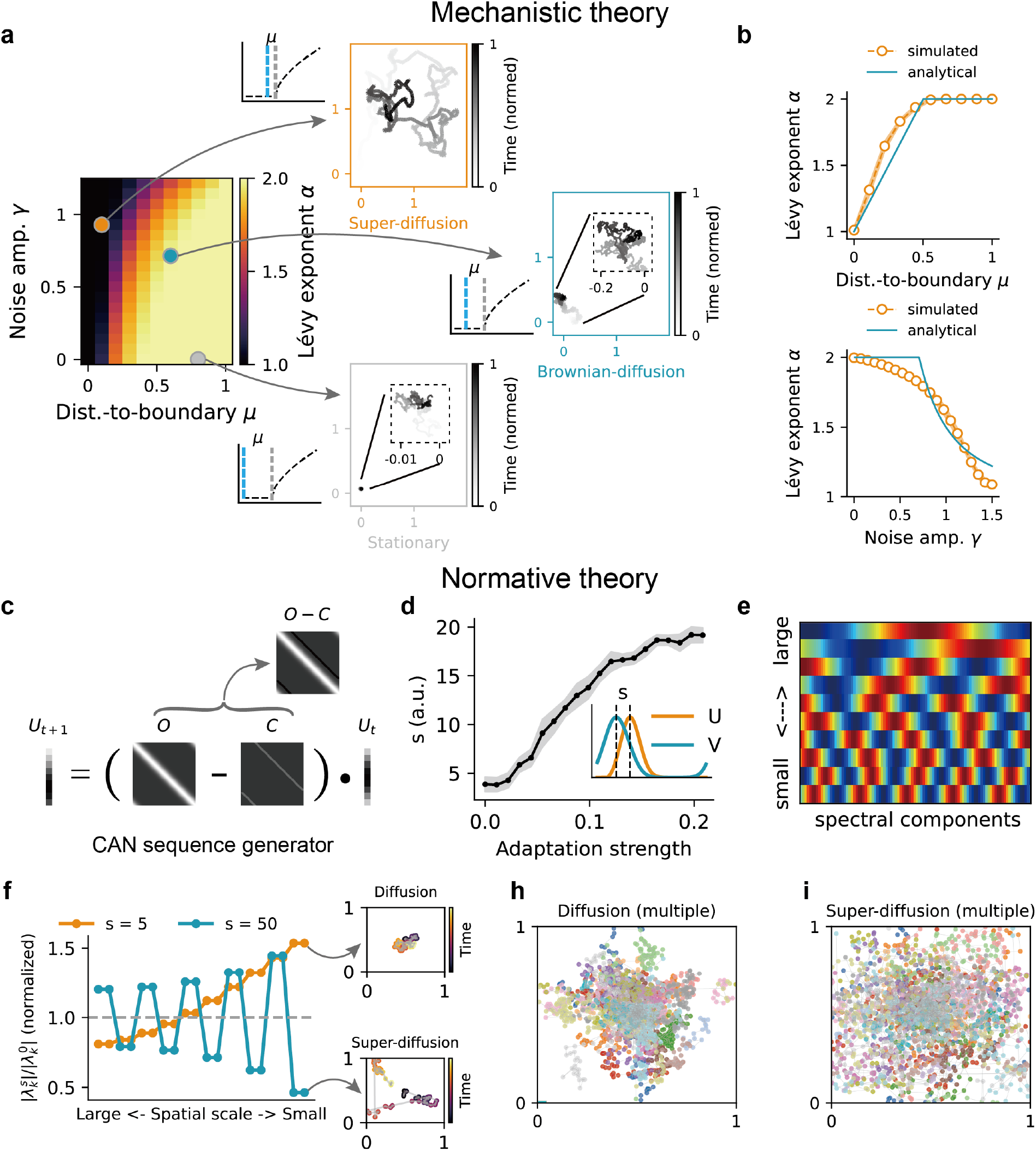
Theory of generating diverse diffusive replay sequences in the CAN model with firing rate adaptation. **(a-b)**, mechanistic theory by solving the network dynamics. **(c-i)**, normative theory by spectral analysis of the state transition matrix. **(a)**, the phase diagram of replay diffusivity as a function of adaptation strength and noise level, measured by the power-law exponent in Eq. (4). Larger values represents super-diffusive dynamics in the generated sequences. The orange, blue and grey dots represent three different parameter settings for super-diffusive, diffusive and stationary dynamics, respectively. The blue dashed lines in three insets represent the value of adaptation strength whereas the grey dashed lines represent the travelling boundary. **(b)**, top panel: the diffusivity of replay sequences as a function of the adaptation strength. Bottom panel: the diffusivity of replay sequences as a function of the noise level. **(c)**, the CAN sequence generator. *U*_*t*_ is the state distribution, *O* is the transition matrix and *C* is the perturbation matrix from the effect of adaptation (negative feedback). **(d)**, Perturbation offset *s* as a function of adaptation strength *m*. Inset: example offset between the activity bump *U* and the adaptation bump *V* . **(e)**, Example eigenvectors (Fourier modes)of the perturbed transition matrix, represented by wave vectors of increasing frequencies (decreasing eigenvalues) (Equation 43). **(f)**, left: rescaled eigenvalues (normalized by original eigenvalues) after the perturbation at different offsets (orange: s=5; blue: s=50; top 20 eigenvalues are shown for better illustration). Right: sampled trajectories from the state sequences based on the perturbed generator (diffusion: s=5; super-diffusion: s=50). **(h)**, multiple diffusion trajectories with s=5. **(i)**, multiple super-diffusion trajectories with s=50.

### 2.2.2 Spectral analysis from a normative viewpoint

The second analysis is treating the CAN model as a sequence generator [McNamee et al., 2021] by vectorizing the dynamics into a master equation with the form of:

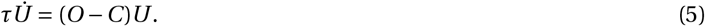

Here, *U* represents the state distribution of the place cell population, *O* represents the transition matrix among states dictated by the recurrent connections, and *C* represents the perturbation matrix modulated by the effect of adaptation. In this scenario, the generation of replay sequences can be viewed as a sampling process from *U*_*t*_, which evolves over time based on the perturbed transition matrix (Fig. 3c). Based on Eq. (5), we can analyze the spectrum of the perturbed transition matrix to gain normative insights about how perturbations lead to specific diffusivity profiles in generated sequences (Fig. 3d-f and Methods. 4.12).

Specifically, the perturbation matrix *C* is a sub-diagonal matrix representing the effect of the adaptation signal *V* on the main activity bump *U* . The adaptation input is of the same profile as *U*, but lags behind *U* with a displacement of *s* (Fig. 3d inset). Stronger adaptation leads to a larger displacement *s* between *V* and *U* (Fig. 3d), resulting in a larger perturbation offset of the matrix *C* (further from the diagonal). Theoretically, the perturbed transition matrix shares the same set of eigenvectors independent of the offset value [Yu et al., 2020, Bracewell, 1986], which are the set of Fourier modes (Fig. 3e) (Methods. 4.12). Therefore, to track the effects of the adaptation-based perturbation on the state distribution over time, we only need to know the perturbation to the spectrum corresponding to perturbed matrices with different diagonal offset values (i.e., different adaptation strengths).

With a small offset (*s* = 5), indicating weak adaptation, the eigenvalues of larger spatial scales (corresponding to lower-frequency Fourier modes in Fig. 3e) are dampened, while those of smaller spatial scales are amplified (Fig. 3f&h). This type of eigenvalue rescaling leads to an increased sampling of local areas, favoring Brownian diffusion. Conversely, with a large offset (*s* = 50), indicating strong adaptation, an increase in eigenvalues of larger spatial scales precedes a decrease in eigenvalues of smaller spatial scales (Fig. 3f&i). This type of eigenvalue rescaling amplifies transition probabilities associated with distant states, favoring super-diffusion. By rigorously deriving the spectrum of the rescaling profile of the eigenvalues under the perturbation along each offset value, we can determine the precise boundary values for the offset at which the sampling patterns switches from Brownian-to super-diffusion (Methods. 4.12 and Fig. S2).

Our spectral analysis provides an insight into the occurrence of diffusive/super-diffusive replay profiles. Importantly, it connects a mechanistic implementation with previous normative theory [McNamee et al., 2021, McNamee, 2024] which treated hippocampal sequence generation as the systematic rescaling of eigenvalues (gain modulation) along the dorsoventral axis of MEC grid cell modules in the hippocampal-entorhinal loop (see the Discussion for more details).

### 2.3 Model predictions

Our model demonstrates that variations in firing rate adaptation could account for the observed diversity of replay diffusivity in experimental data. Building on these findings, our model posits three novel predictions (Results. 2.3.1-2.3.3), each supported by our empirical analysis of experimental data (Results. 2.4.1-2.4.3). By relating replay diffusivity to theta sequences across animals, distinguishing between different brain states within individual animals, and accounting for the relationship between neural activity and replay step size within individual replay sequences, these results enrich our understanding of the functional roles of replay dynamics at multiple hierarchical levels.

#### 2.3.1 Replay diffusivity correlates with theta sequence length

It is well-established that when animals navigate an environment, hippocampal sequences exhibit theta sweeps, wherein decoded positions locally oscillate around the animal’s location at theta frequency [Foster and Wilson, 2007]. Conversely, when animals are stationary, hippocampal sequences display non-local sequential activity, indicating positions away from the animal’s location [Lee and Wilson, 2002]. While theta sequences are posited to contribute to the formation of initial memory traces and replay sequences to the consolidation of these memories [Drieu et al., 2018], the intricate dynamic relationship connecting these two phenomena remains underexplored. Here we predict a correlation between more diffusive replay sequences and longer theta sequences (Fig. 4a), positing that an increase in firing rate adaptation strength underpins the augmentation of both types of sequential dynamics. To generate theta sweeps in the model, we drew on previous work [Chu et al., 2024] by introducing location-dependent sensory input (modeled as an external bump input) to the CAN. The interplay between this external bump input and the intrinsic dynamics from firing rate adaptation facilitates bump sweeps around the animal’s location when it is moving in the environment (see Methods. 4.8 for details). With an increase in adaptation strength, the activity bump sweeps further away from the animal’s location (Fig. 4a). Conversely, when sensory input is absent (mimicking the resting state), replay sequences emerge and exhibit increased diffusivity with the adaptation strength. Integrating modeling results for both replay and theta sequences, our model predicts a positive correlation between replay diffusivity and theta sweep length (Fig. 4; Pearson correlation *r* = 0.77, *p* < 0.001), underscoring that both phenomena are modulated by the strength of firing rate adaptation (see Results. 2.4.1 for the experimental data analysis).

**Figure 4:**
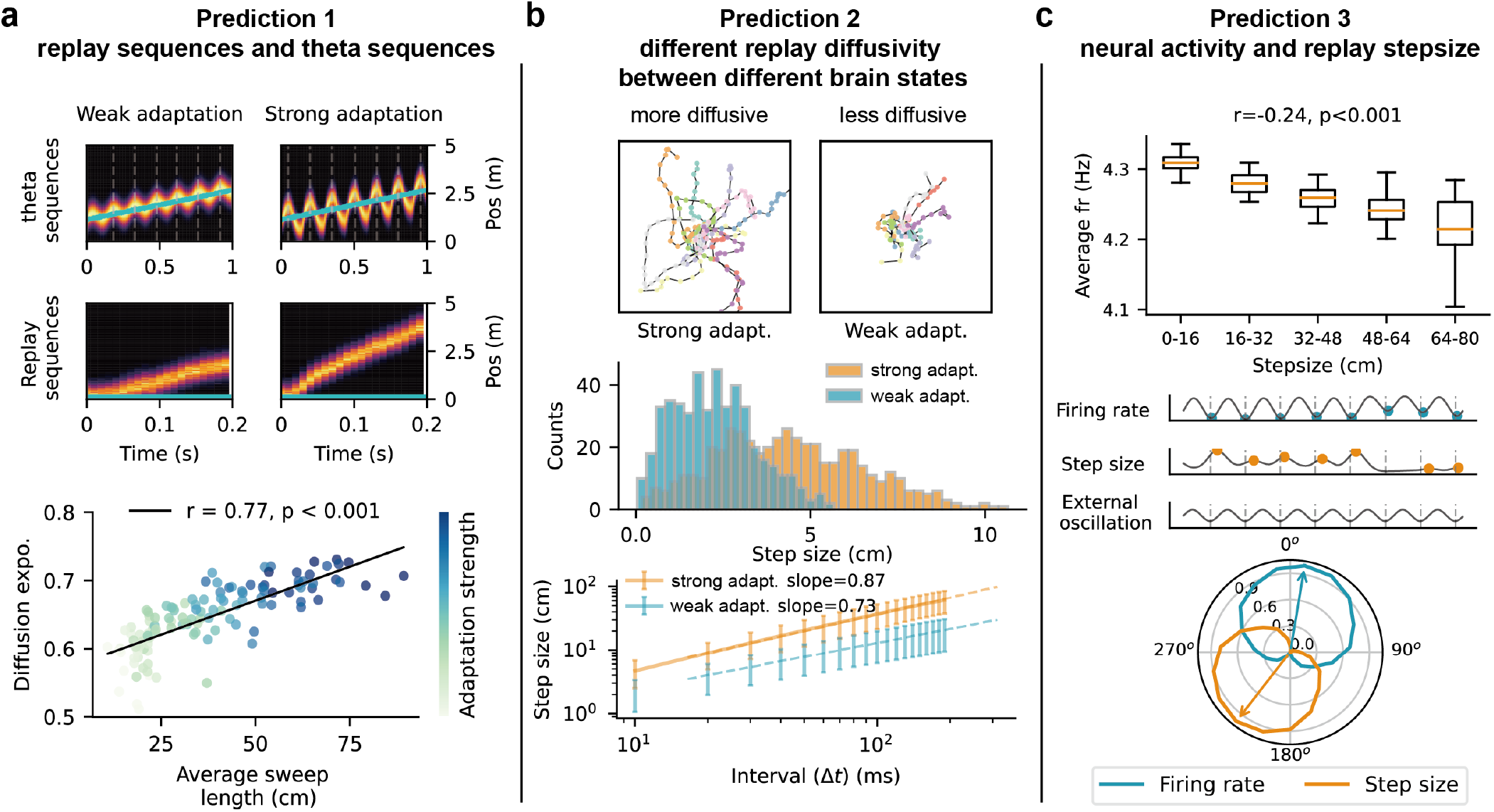
Model predictions. **(a)**, we predict that across animals, more diffusive replay sequences correlate with longer theta sequences (both reflecting stronger adaptation). Top panel: Theta sequences (top row) and replay sequences (bottom row) in the CAN model under conditions of weak and strong firing rate adaptation, respectively. Blue lines depict the animal’s trajectories, while heat maps illustrate network activity. Bottom panel: a scatter plot showing the positive correlation between the diffusivity of replay sequences and the length of theta sequences. Each dot represents a simulated CAN network with varying adaptation strength, with darker blue indicating stronger adaptation. **(b)**, we predict that within an animal, firing rate adaptation varies with different brain states, which leads to a difference in replay diffusivity. Top panel: example trajectories (depicted in different colors) under conditions of strong and weak adaptation, corresponding to more and less diffusive replay dynamics, respectively. Middle panel: a histogram comparing the step sizes for the two trajectory types shown in the top panel, illustrating a heavier-tailed distribution under strong adaptation strength. Bottom panel: a plot of step size versus time duration on a log-log scale highlights a larger slope for simulated replay trajectories (orange) compared to sleep replay trajectories (blue), with dashed lines indicating linear fits. **(c)**, we predict that within a single replay trajectory, increases in neural excitability, in cooperation with firing rate adaptation, reduces the step-size within individual replay sequences. Top panel: negative correlation between the mean firing rate and replay step size, where longer step sizes are associated with lower firing rates in the network. Middle panel: both the firing rate (top) and step size (middle) are phase-locked to an external slow-gamma oscillation (bottom), with blue and orange dots highlighting the troughs and peaks in the firing rate and step size curves, respectively. Dashed lines indicate the troughs of the slow-gamma oscillation. Bottom panel: normalized contour plots and circular weighted mean (arrows) for neural activity (blue) and step sizes (orange) as a function of the slow-gamma phase.

#### 2.3.2 Different replay diffusivity between different brain states

It has been shown that neurotransmitter levels such as acetylcholine, dopamine, and noradrenaline differ across different brain states [Samanta et al., 2020]. We predict that the adaptation strength may change across different brain states [Yoshida et al., 2013, Barkai and Hasselmo, 1994, Madison and Nicoll, 1984], which further leads to a difference in replay dynamics between different behavioural states within individual animals (Fig. 4b). Specifically, increased adaptation strength leads to super-diffusive dynamics characterized by a heavy-tailed power-law distribution of step sizes (Fig. 4b). Conversely, decreased adaptation strength results in less diffusive dynamics, indicated by a Gaussian distribution of step sizes (Fig. 4b). We quantitatively differentiate these dynamics using the diffusion exponent, calculated as the slope of step size against time duration on a log-log scale (Fig. 4b). We use this measurement because it has been widely used in previous experimental literature [Stella et al., 2019, McNamee et al., 2021, Krause and Drugowitsch, 2022], and it correlates with the power-law exponent from the theoretical analysis (Eq. (4), see Methods. 4.9 for more details). We show that in the model, a super-diffusive sequence exhibits a diffusion exponent greater than 0.5, while a diffusive sequence features an exponent approximately 0.5. Although we lack direct evidence of the variance in adaptation strength across behavioural states (see Discussion for more details), these modelling results motivate us to examine if there is a difference in replay dynamics during awake and sleep states within individual animals (Results. 2.4.2).

#### 2.3.3 Inverse relation between population activity and replay step size

It has been observed that individual awake replay sequences exhibit interleaved local- and long-jump movements [Pfeiffer and Foster, 2015]. However, how the detailed structure of replay step sizes is modulated remains largely unknown. We predict that neural activity negatively correlates with step sizes within individual replay sequences. Specifically, increase in neural excitability counterbalances the effect of adaptation, leading to smaller replay step sizes, while decrease in neural excitability provides less counterbalance, resulting in larger replay step sizes. Consequently, this leads to a negative correlation between population activity and replay step size (*r* = −0.24, *p* < 0.001, Pearson correlation; Fig. 4c). Furthermore, by incorporating neural oscillations as external input in our model (typically slow-gamma oscillations during replay events, mimicking those generated in CA3; see Methods. 4.11 for more details), we provide an explanation for previous experimental findings [Pfeiffer and Foster, 2015] that showed replay step size and neural activity are modulated by the slow-gamma rhythm (25-50 Hz). Specifically, the preferred phase of large replay step size opposes that of neural activity: large replay step sizes occur at the trough of the slow-gamma cycle, where neural activity is low, while local movements (small step sizes) align with the peak, where neural activity is high (Fig. 4c, also see Results. 2.4.3 for more details).

### 2.4 Verifying model predictions with experimental data

Here we verify these model predictions in empirical data which include tetrode recordings from the hippocampi of 9 rats performing a 15-min spatial alternation task on a W-shaped track [Karlsson et al., 2015] (Fig. S1; see Methods. 4.1 for more details). To decode theta sequences and replay sequences, we employed a “clusterless” state space decoding algorithm [Denovellis et al., 2021] (see Methods. 4.2 for more details). This decoder reliably estimated the animal’s position (with a median error of 9.0 cm, and 95% CI of 8.2 −9.8 cm), comparable to results from prior studies [Davidson et al., 2009, Farooq and Dragoi, 2019]. For theta sequences, we segmented the running periods (where the animals’ speed exceeded 4 cm/s) into five folds. We trained the model on four folds and decoded theta sequences on the remaining one (see Fig. 5b&c and Fig. S3 for more examples). For replay sequences during the task-engaged immobile state, we trained the model on the running data and applied it to SWR events (172 ± 62 SWRs per recording epoch) (Fig. 5b&c).

**Figure 5:**
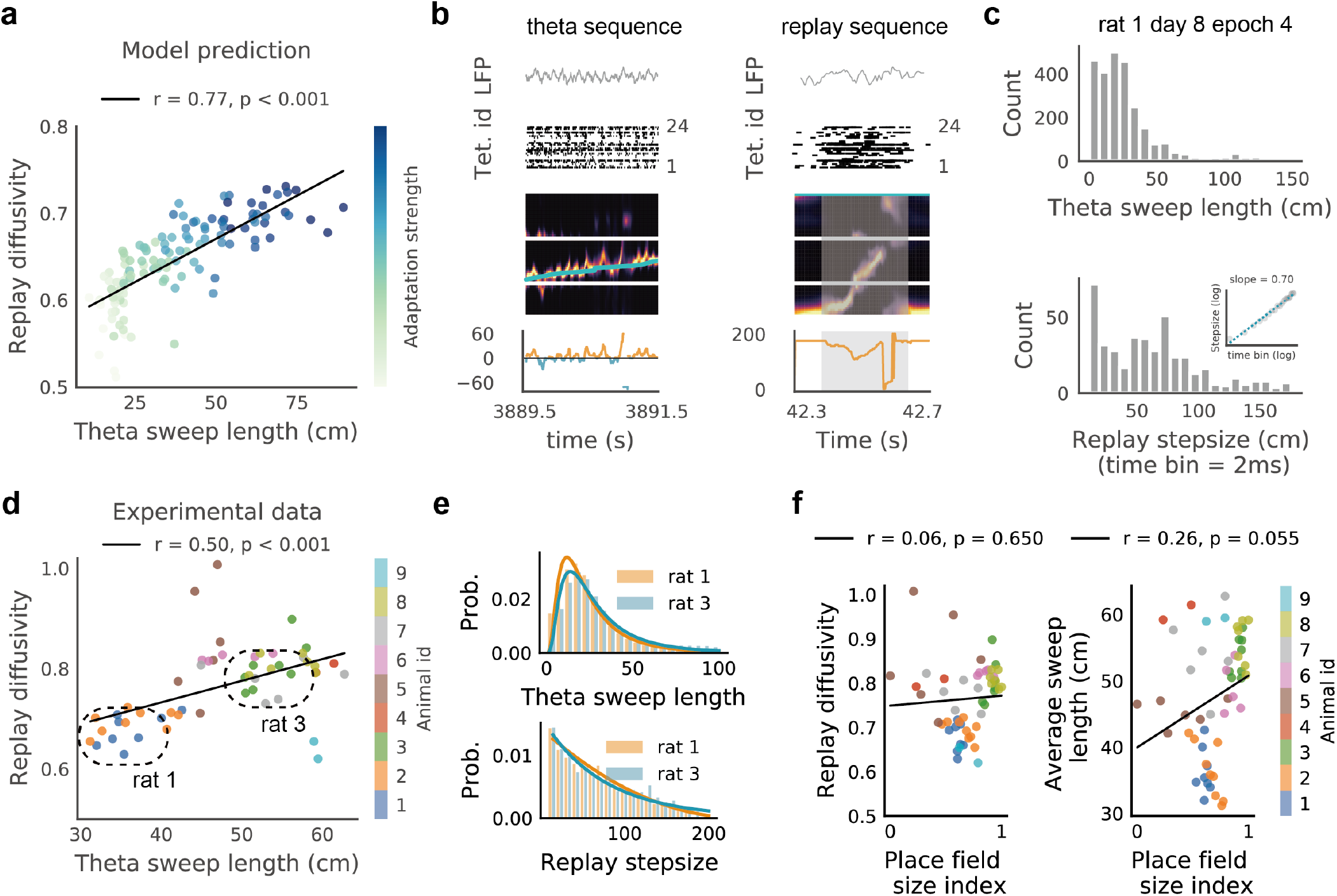
More diffusive replay sequences are associated with longer theta sequences. **(a)**, model prediction (refer to Fig. 4a). **(b)**, an illustrative example of a theta sequence and a replay sequence in the dataset. From top to bottom in each panel: LFP signal from a CA1 tetrode; multiunit activity; the posterior probability map; offset distance as a function of time (on the left: blue represents look-back distance, and orange represents look-ahead distance; on the right: offset distance between the decoded position and the actual position). **(c)**, histograms of theta sequence length (from all LFP theta cycles) and awake replay step size (time bin is 2 ms) for one recording epoch from Rat 1. The inset marks the relationship between replay step size and the time bin on the log-log scale. **(d)**, correlation between the diffusivity of replay sequences and the length of theta sequences in the real data. Each point represents the diffusion exponent and the average length of theta sequences from one recording day, with different colors denoting data from various animals. Dashed circles highlight the data from two exemplar rats shown in **(e). (e)**, top panel: distribution of theta sequence lengths for Rat 1 and Rat 3 with log-normal curve fitting, noting that Rat 3 exhibits longer theta sequences. Bottom panel: distribution of replay step sizes for Rat 1 and Rat 3 with generalized Pareto curve fitting, highlighting that Rat 3 shows a longer-tailed distribution of replay step size. **(f)**, left panel: diffusion exponent as a function of the place field size index, with different colors representing data from various animals. Right panel: length of theta sequences as a function of the place field size index.

#### 2.4.1 Replay diffusivity correlates with theta sequence length

Our model predicts a positive correlation between the length of theta sequences and the diffusivity of replay sequences across animals (Fig. 5a). In the empirical data, we also discovered a significant correlation between awake replay diffusivity and average theta sweep length (Fig. 5d; Pearson correlation *r* = 0.50, *p* < 0.001), indicating that more diffusive awake replay is associated with longer theta sweeps during active running (see Fig. 5e for the data from two rats). While the correlation between replay diffusivity and theta sequence length is evident in the aggregated data from all animals, individual plots for each animal also reveal a trend of correlation between these two variables (Fig. S4). To ascertain that this correlation was not coincidental, we correlated theta sweep lengths with replay diffusivities randomly selected from other recording days within the same animal, resulting in significantly lower correlation coefficients (Fig. S5; p=0.014, Kolmogorov-Smirnov test). Additionally, we explored the correlation between theta sequence length and replay diffusivity during subsequent sleep periods, and found that the positive correlation still held (Fig. S6; Pearson correlation *r* = 0.49, *p* = 0.005).

It is possible that the place field size might act as a confounding factor to the observed correlation between replay diffusivity and theta sweep length. Specifically, smaller place fields could prevent the network activity from spreading away, resulting in shorter theta sweep length and less diffusive replay sequences. To exclude this potential confound, we calculated the place field size index using the population vector correlation method [Battaglia et al., 2004] (see Methods. 4.10 and Fig. S7). Our analysis showed a marginal correlation between place field size and theta sweep length (Pearson correlation *r* = 0.26, *p* = 0.055), indicating that larger place fields are associated with longer theta sequences. However, there was no significant correlation between place field size and replay diffusivity (Pearson correlation *p* = 0.650; Fig. 5f), indicating that place field size is not a major confounding factor for replay diffusivity. Additionally, we excluded decoding accuracy as a potential confounding factor. Specifically, a recording session with fewer spike events might result in noisier decoding for both theta and replay sequences, which could artificially lengthen theta sequences and increase replay diffusivity. However, the correlation between theta sweep length and the number of spikes involved in decoding was weak (Pearson correlation *r* = 0.31, *p* = 0.023), and there was no significant correlation between replay diffusivity and the number of spikes (Pearson correlation *r* = 0.14, *p* = 0.317) (Fig. S8). This indicates that the observed correlation between replay diffusivity and theta sequence length is not substantially influenced by variations in decoding accuracy.

In summary, our analysis quantitatively reveals a relationship between the length of theta sequences and the diffusivity of awake replay, underscoring their coordinated role in hippocampal cognitive functions, especially memory formation and consolidation [Drieu et al., 2018, Muessig et al., 2019]. This relationship was predicted by our model as they are both affected by the strength of firing rate adaptation (see Discussion for more details).

#### 2.4.2 Different replay diffusivity between awake and sleep states

We have demonstrated that animals exhibiting more diffusive replay tend to have longer theta sequences during the same task-engaged phase, a phenomenon observed both in our model and the neural data. However, whether replay dynamics vary in the same hippocampal network remains unclear. Previous studies have shown that awake replay is super-diffusive when animals performed a spatial memory task [Pfeiffer and Foster, 2015, Krause and Drugowitsch, 2022], while sleep replay is Brownian-diffusive after animals performed a random foraging task [Stella et al., 2019]. While these findings are based on different animals engaged in different tasks, it remains unclear whether such a difference holds in the same animal with different behavioural states, or it is simply due to the difference of the engaged tasks (the spatial memory task versus the random foraging task). Our model suggests that varying the adaptation strength leads to a difference in replay dynamics in the simulated hippocampal network without changing other network configurations (Fig.6a). Inspired by this prediction, we compared replay diffusivity during both task-engaged immobile state and subsequent sleep state for the same animal. We found that task-engaged awake replay exhibits significant super-diffusive dynamics (Fig.6b; diffusivity with 0.74 ± 0.15; Wilcoxon signed-rank test with *p* < 0.001), aligning with prior research [Pfeiffer and Foster, 2015, Krause and Drugowitsch, 2022]. To ensure these super-diffusive dynamics were not due to random ordering of spike events, we performed a shuffle control [Stella et al., 2019] by randomizing the decoded positions within trajectories (equivalent to shuffling time bins while preserving a constant set of decoded locations in space). This control yielded diffusivity values near zero, significantly lower than the 5_*th*_ percentile of the actual diffusivity values during awake replay (Fig.6b).

To identify sleep replay, we extracted candidate sleep periods in the resting box as non-REM phases with high-amplitude LFP (LIA) periods, utilizing LFPs from CA1, CA2, and CA3 tetrodes (see Fig. 6c and Methods. 4.1 for more details). We included only SWRs from these candidate sleep periods for further analysis (248 ± 154 SWRS per recording epoch). Subsequently, we decoded sleep replay trajectories using the place fields measured in the preceding running session (Fig. 6d) and calculated one diffusion exponent for each sleep session (Fig. 6e). The analysis revealed that sleep replay is significantly less diffusive than the awake replay dynamics that preceded it (53 paired awake-sleep recording epochs in total; Wilcoxon signed-rank test with *p* < 0.001) (Fig. 6f). It is noteworthy that the diffusivity values of sleep replay in our study are significantly larger than 0.5 (diffusivity with 0.69 ± 0.12; Wilcoxon signed-rank test with *p* < 0.001), diverging from the Brownian-diffusive dynamics reported in previous research [Stella et al., 2019] (Fig. S9). This discrepancy may stem from differences in task design: the spatial alternation task on a W-shaped track constrains the range of random exploration, in contrast to the open field of the random foraging task, which permits movement in multiple directions.

**Figure 6:**
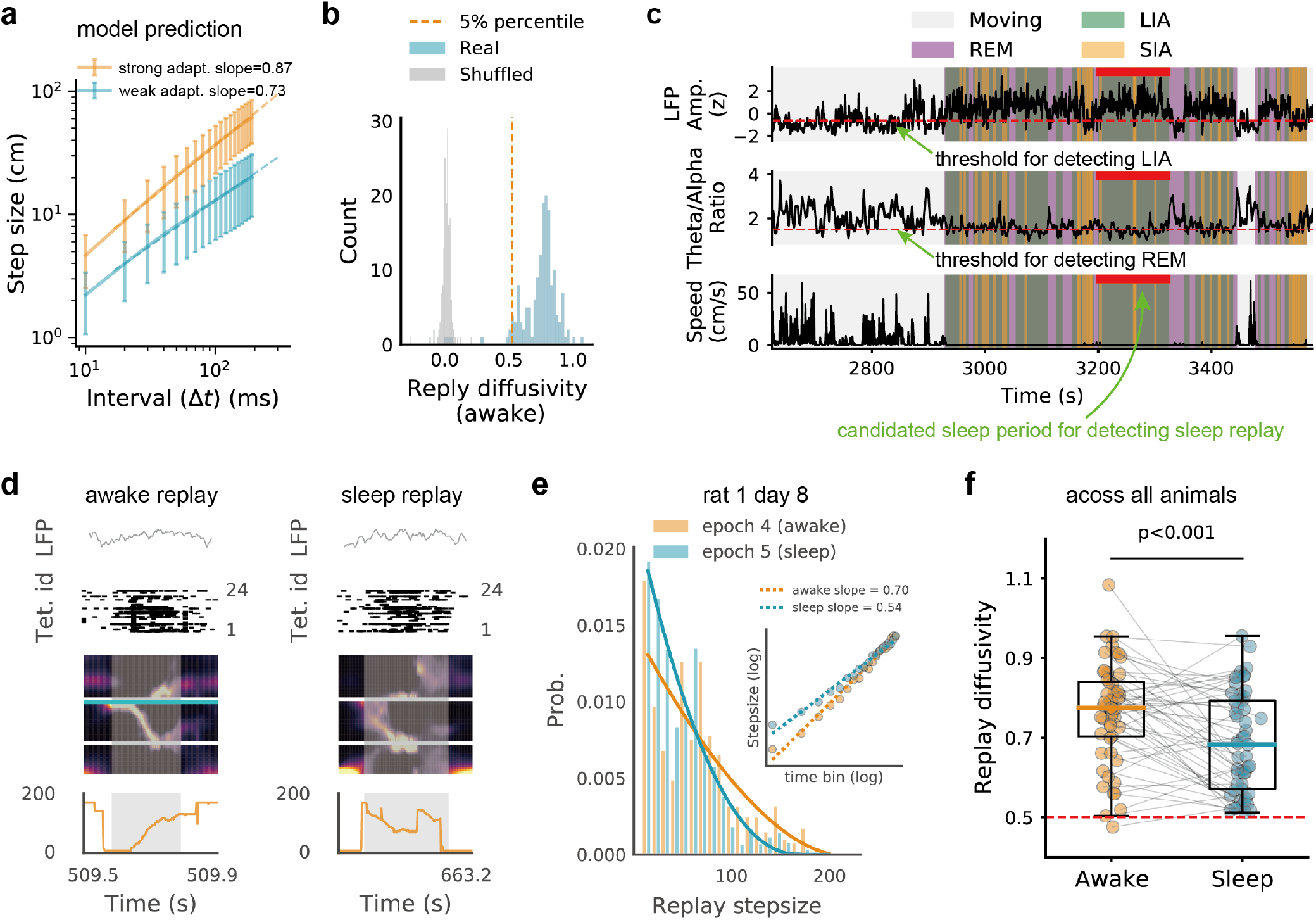
Replay sequences exhibit greater diffusivity in the awake immobile state compared to the subsequent sleep state. **(a)**, model prediction (refer to Fig. 4b). **(b)**, histogram of diffusion exponents (one per running session) for awake replay across all sessions and animals (in blue) and diffusion exponents from shuffled position order (in grey). The orange dashed line indicates the 5th percentile of the measured diffusivity values. Note that the awake replay is significantly larger than 0.5 and hence super-diffusive. **(c)**, sleep state identification as the animal in the resting box. Top, aggregated LFP amplitude from CA1, CA2, and CA3 tetrodes for SIA/LIA period detection; middle, theta to delta ratio for REM period detection; bottom, running speed for immobile period detection. Red bars mark candidate sleep periods lasting at least 90 s with extended (>5 s) continuous LIA periods (see Methods.4.1). Only ripple events within these periods were analyzed further. **(d)**, examples of an awake replay sequence (left) when an animal performed the W-track spatial alternation task and a subsequent sleep replay sequence (right) when the animal was in the resting box (refer to Fig. 5 for panel descriptions). **(e)**, step size distribution (time bin = 2 ms) for awake (orange) and subsequent sleep (blue) replays from two successive sessions for one animal. The inset illustrates step size versus time bins on a log-log scale for both replay types, showing greater diffusivity in awake replay. **(f)**, comparison of diffusivity of awake replay (orange) and subsequent sleep replay (blue) across all recording sessions and animals. Each dot represents a diffusion exponent from a recording session, with grey lines connecting values from successive running and sleep sessions. Awake replays show significantly higher diffusivity than sleep replays (Wilcoxon signed-rank test, *P* = 4.6 × 10^−4^).

This change in replay dynamics highlights distinct information processing mechanisms during different resting states. Specifically, replay sequences in the task-engaged immobile state may reflect the animal’s recent movements [Krause and Drugowitsch, 2022] and are potentially linked to planning and decision-making processes [Pfeiffer and Foster, 2013]. In contrast, sleep replay, which is less directly tied to recent experiences, may support generalization across various behavioral outcomes [Stella et al., 2019]. While the mechanism that allows individual hippocampal networks to switch between these replay dynamics during awake and sleep states is still lacking, our model suggests that such alternation could be achieved by simply adjusting the adaptation strength (see Discussion for more details).

#### 2.4.3 Inverse relation between population activity and replay step size

We have demonstrated that firing rate adaptation can modulate replay diffusivity across different animals and brain states within an individual animal. However, it remains unclear whether this modulation occurs within individual replay events. Our model predicts a negative correlation between neural activity and replay step size (Fig. 7a; *r* = −0.24, *p* < 0.001, Pearson correlation), suggesting that increased neural excitability counterbalances the effect of adaptation, leading to smaller replay step sizes. To validate this prediction with empirical data, we analyzed replay events across all animals during the awake immobile state, revealing a significant negative correlation between replay step size and the aggregate spike count.(Fig.7b; *r* = −0.12, *p* < 0.001, Pearson correlation). Specifically, during periods of high neural activity, spatial representation tends to focus on a single location, whereas during periods of reduced neural activity, spatial representation is more prone to transition to different locations. Although decreased activity (fewer spikes in the decoding time window) might affect decoding accuracy and potentially inflate replay step size, our model offers a theoretical basis that elucidates and substantiates this empirical finding.

**Figure 7:**
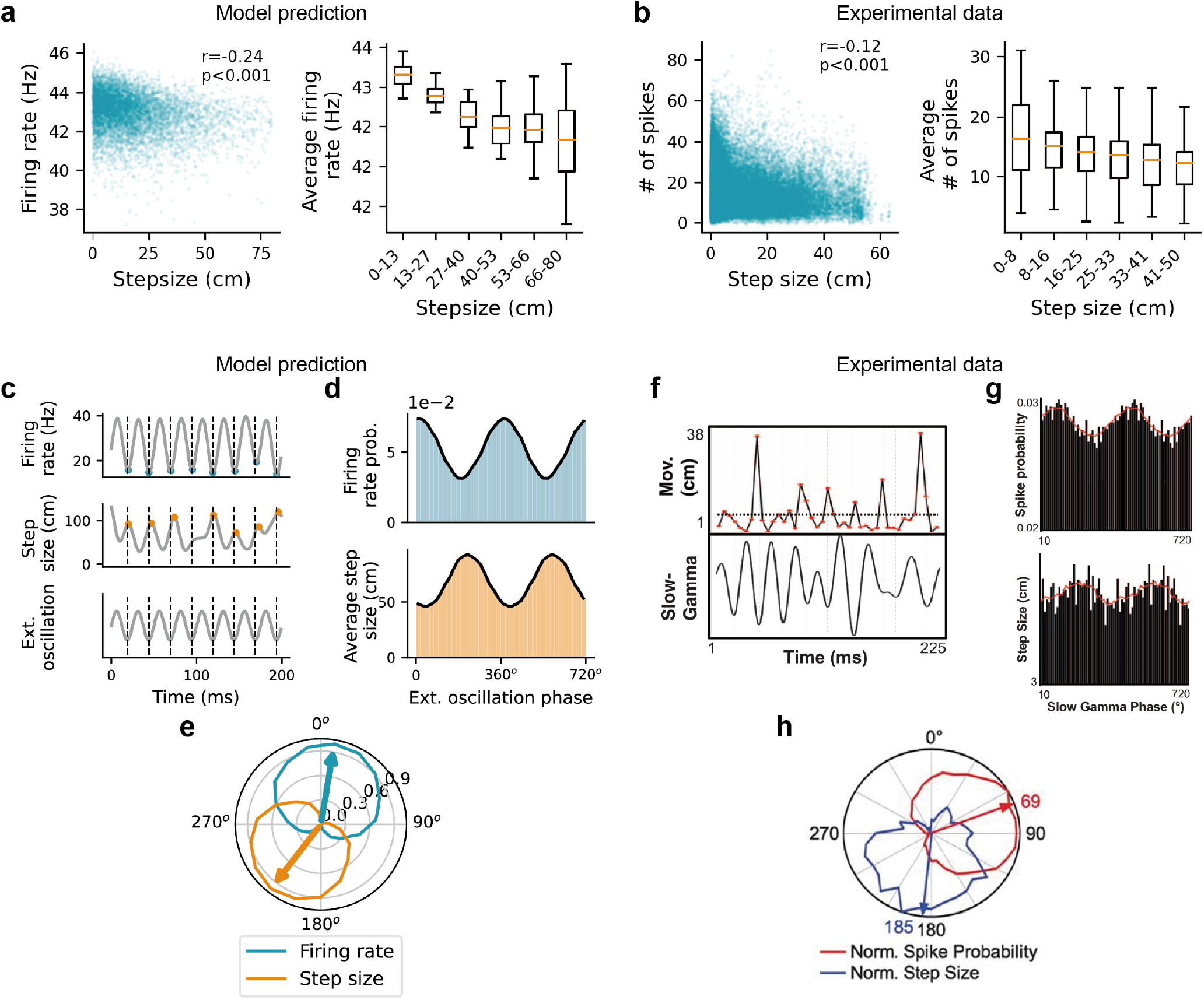
Inverse relation between population activity and replay step size, and their relationships to slow-gamma oscillations. **(a)**, model prediction showing the negative correlation between neural activity and replay step size (refer to Fig. 4c). Left panel: scatter plot of the population firing rate (summed over all neurons within the time window during which the step size is calculated) against the step size. Each dot represents a sample from the replay trajectories simulated with varying adaptation strengths. Right panel: box plot of the population firing rate versus binned step size from the left panel. Orange lines indicate mean values, boxes represent the quartile range, and bars show non-outlier range extremes. **(b)**, empirical observation of the negative correlation between replay step size (time bin = 20*ms*) and neural activity in the spatial alternation task. **(c-e)**, model prediction of anti-phase locking between neural activity and replay step size to slow-gamma oscillations. **(c)**, phase locking of population firing rate (top) and replay step size (middle) with an external slow-gamma oscillation (bottom). Blue and orange dots highlight troughs and peaks in the firing rate and step size curves, respectively. Dashed lines mark the troughs of the slow-gamma oscillation. **(d)**, firing rate probability (top) and step sizes (bottom) as a function of the slow-gamma phase (displaying two repeated cycles). **(e)**, normalized contour plots and circular weighted mean (arrows) for neural activity (blue) and step sizes (orange) as a function of the slow-gamma phase. **(f-h)**, slow-gamma phase-locking phenomenon of neural activity and replay step size, adapted from [Pfeiffer and Foster, 2015] with permission. **(f)**, movement (top) and slow-gamma oscillation (bottom) for a representative trajectory event from previous experimental data [Pfeiffer and Foster, 2015]. **(g)**, across all open field running sessions, the aggregate spike probability (top) and step size (bottom) as functions of the slow-gamma phase (bin size = 10^°^) for all trajectory events. Red lines indicate the running box average (box size = 8 bins). **(h)**, normalized contour plots with circular weighted mean (arrow) for box-average spike probability (red) and step size (blue) relative to the slow-gamma phase.

Building upon the observed negative correlation, our model further predicts that neural activity and replay step size are anti-phase locked to external neural oscillations (Fig. 7c-e; see Methods. 4.11 for more details). Specifically, at the peaks of these oscillations (corresponding to maximal excitatory input), population activity is high. This elevated activity counterbalances the negative feedback effect from firing rate adaptation, resulting in small replay step size. Conversely, at the oscillation troughs (minimal excitatory input), population activity decreases, which minimizes the counterbalancing effect and allows the activity bump to transition between locations, leading to larger replay step size. Intriguingly, when oscillations align with the slow-gamma rhythm, which intensifies during replay events, our model explains previous empirical findings [Pfeiffer and Foster, 2015] where they showed the preferred phase of large step size locked to the slow-gamma rhythm (Fig. 7f). Moreover, the preferred phase of neural activity is inversely related to that of large replay step sizes (Fig. 7g&h). These results suggest that firing rate adaptation and neural excitability can cooperatively modulate replay dynamics, shaping the temporal structure of replay sequences in a more detailed way.

## 3 Discussion

In this study, we presented a theoretical framework to capture the diverse diffusivity of replay dynamics in the hippocampus by modeling the place cell population as a continuous attractor network (CAN) with firing rate adaptation (Results. 2.1). Firing rate adaptation destabilizes the activity bump, inducing intrinsic mobility within the network. This intrinsic mobility interacts with internal network noise, resulting in a spectrum of replay sequences exhibiting different diffusivities. Through both theoretical and empirical analysis, our model reconciles a range of previously observed, seemingly disparate replay dynamics, including stationary, Brownian, and super-diffusive dynamics(Results. 2.2). Furthermore, we made predictions at three levels—across animals, across brain states within animals, and across individual replay sequences—and verified them with empirical data. First, we demonstrated a positive correlation between the diffusivity of replay sequences and the sweep length of theta sequences, both reflecting stronger adaptation (Results. 2.3.1&2.4.1). Second, we found that the diffusivity of awake replay is significantly higher than that of subsequent sleep replay within individual animals (Results. 2.3.2&2.4.2). Third, at the level of individual replay sequences, we revealed a detailed structure of replay dynamics: replay step sizes negatively correlate with population activity, which also explains the slow-gamma modulation of replay step size observed previously (Results. 2.3.3&2.4.3). These findings suggest that flexible modulation of firing rate adaptation plays a crucial role in the dynamics of hippocampal sequences, raising several issues concerning hippocampal functions and theoretical modeling, which are addressed below.

There are two main types of computational models explaining hippocampal sequence generation. The first type focuses on internal hippocampal circuit dynamics, incorporating mechanisms such as asymmetric synaptic connections [Tsodyks et al., 1996], cholinergic modulation [Hasselmo et al., 1995, Saravanan et al., 2015], firing rate adaptation [Hopfield, 2010, Itskov et al., 2011, Azizi et al., 2013] and short-term plasticity [Romani and Tsodyks, 2015, Pietras et al., 2022, Singh et al., 2022]. Despite differences in their biological implementations, these mechanisms rely on the symmetry-breaking effect in network dynamics, temporarily destabilizing the activity bump and introducing autonomous transitions among network states. The second type focuses on the entorhinal-hippocampal system (EHC model) [McNamee et al., 2021], which forms a linear feedback network between the two brain regions. In this model, spectral modulation along the dorsoventral axis of MEC grid cells modulates the generative sampling process of state sequences in the hippocampus. This idea is supported by empirical data showing that MEC input controls the temporal organization of hippocampal activity [Schlesiger et al., 2015, Yamamoto and Tonegawa, 2017]. While our model’s sequence generation does not depend on systematic modulation of the MEC grid population, it connects to the EHC model in several ways. Both models utilize feedback modulation to recirculate the update of state sequences. In the EHC model, this involves slow dynamics in the large spatial loop between the two brain regions, while in our model, it involves slow dynamics in the large temporal loop due to firing rate adaptation. Additionally, for generating distinct sequences, the EHC model varies the power spectrum of the generator, akin to different gain control across grid modules, while our model varies the adaptation strength. This difference can be reconciled by a spectral analysis of CAN dynamics, revealing a different form of spectral modulation in generating hippocampal sequences from the EHC model (see Fig. 3 and Fig. S2). Computationally, these mechanisms are not mutually exclusive. Instead, they share a common form of slow feedback modulation, exemplified as firing rate adaptation in our study. Indeed, these mechanisms may cooperate to offer a more comprehensive explanation of sequence generation in the hippocampus.

The coordination between theta sequences and replay sequences (Fig. 5e) aligns with their co-development in the hippocampus. Specifically, replay events are stationary in pre-weaning animals and become more sequential after weaning, corresponding with the gradual maturation of theta sequences [Muessig et al., 2019]. This development might be attributed to the gradual maturation of attractor dynamics in the hippocampus [Wills et al., 2005], supported by the finding of increased co-firing between cells with overlapping place fields during SWRs [Muessig et al., 2019]. In our model, a more mature attractor network corresponds to stronger connections among neurons, within which firing rate adaptation triggers the activity bump to move more smoothly, leading to more diffusive replay sequences and longer theta sweeps. Previous experimental work also revealed a causal link between theta sequences and replay sequences, showing degraded replay sequences after perturbing theta sequences [Drieu et al., 2018]. While our theoretical and empirical analysis does not directly establish a causal link between them, we suggest that perturbing theta sequences may weaken the network connectivity through Hebbian plasticity [Jensen and Lisman, 1996, Skaggs et al., 1996, Sato and Yamaguchi, 2003]. This weakened connectivity could prevent the spreading of activity bump due to adaptation, and therefore explains the causal relationship between theta sequences and replay sequences.

Furthermore, our results offer an explanation for how the hippocampus adopts different strategies of sequence generation during awake and sleep SWRs. The higher diffusivity of awake replay compared to sleep replay, which closely resembles the diffusivity of behavioral trajectories, suggests that awake replay might directly reflect the animal’s movements. This mirroring could be crucial for memory consolidation [Diba and Buzsáki, 2007, Krause and Drugowitsch, 2022] and goal-directed planning [Pfeiffer and Foster, 2013]. In contrast, sleep replay seems to focus on generalizing across various behavioral experiences stored in multiple maps, potentially aiding in the formation of novel cognitive maps [Van de Ven et al., 2016, Stella et al., 2019]. Our modeling results suggest that this difference can be modulated by controlling the strength of firing rate adaptation during awake and sleep states. Although our empirical data supports this hypothesis, we acknowledge that the direct examination of adaptation strength differences between awake and sleep states has not yet been achieved. Evidence affecting firing rate adaptation has been derived from in vitro whole-cell patch clamp recordings, where neuromodulators, such as cholinergic modulation, have been shown to affect adaptation strength during awake and sleep states [Madison and Nicoll, 1984, Barkai and Hasselmo, 1994]. However, future research highlights the need for more refined experimental techniques that can directly measure and manipulate adaptation strength in vivo during different SWR-related awake and sleep states. Future studies employing advanced electrophysiological and imaging techniques, as well as genetic and pharmacological manipulations, could provide the necessary data to rigorously test this aspect of our model and experimentally validate our hypothesis.

The simulated theta and replay sequences in our model depend on the translation-invariant form of synaptic connections among neurons. This type of synaptic connection is essential to form a continuous attractor state that allows smooth transitions between attractor states. However, it remains unclear how continuous attractor dynamics are formed in the hippocampal place cell assembly. While we tested the model predictions with empirical data from well-trained animals, where attractor dynamics might be stabilized during repeated tasks, it would be interesting to investigate the formation of attractor dynamics as well. We propose that attractor dynamics and hippocampal theta/replay sequences might co-develop during the learning of environmental structures. Both theta and replay sequences may contribute to the modification of synaptic connections between place cells, aiding in the stabilization of attractor dynamics. Conversely, the stabilized attractor dynamics generate useful sequential representations for various cognitive functions, including exploration, consolidation, and planning. In conclusion, our theoretical framework provides a robust model for understanding the diverse diffusivity of hippocampal replay sequences. By incorporating firing rate adaptation into a continuous attractor network, we explain the mechanisms underlying different replay dynamics observed empirically. This work not only bridges the gap between theoretical predictions and experimental data but also opens new avenues for exploring how intrinsic and extrinsic factors modulate hippocampal functions.

## 4 Methods

### 4.1 Experimental data and SWR detection

#### 4.1.1 Experimental data

The dataset used in this study has been described in details in Karlsson et al. [2015]. On each experimental day, each animal (9 male Long Evans rats in total) performed a 15-min running epoch on a W-shaped track, flanked by 20-min rest sessions in a rest box. The W-shaped track had one reward site at the endpoint of each arm, and the animals were rewarded for performing a continuous alternation task (Fig. S1) [Karlsson and Frank, 2008]. Neural recordings were obtained via a microdrive array containing 30 independently movable tetrodes targeting CA1, CA2, CA3, MEC, Subiculum, and DG, depending on the animal. For analysis in the current study, we only included tetrodes in CA1, CA2, and CA3. Each running epoch consisted of 10-24 tetrodes (140 valid epochs in total; 2.6 ± 0.7 epochs per day, 5.9 ± 2.4 recording days per animal). Multiunit spikes were then obtained as any potential exceeding a 60 *µ*V threshold on any one of the four tetrode wires. The waveform was identified as the electrical potential value on each wire of the tetrode at the time of maximum potential of any of the four wires. All interneurons were excluded from analysis when the spike widths were less than 0.3 ms based on the waveform feature.

#### 4.1.2 Identifying SWRs during awake

Detection of sharp wave and ripple events (SWRs) was performed only when at least three CA1 cell layer recordings were available, following the method described in Kay et al. [2016]. Specifically, LFPs from all available CA1 cell layer tetrodes were band-pass filtered between 150–250 Hz, then squared and summed across tetrodes. This sum forms a single population trace over time, which was then smoothed with a Gaussian kernel (*σ* = 4 ms) and square-rooted. Candidate SWR times were detected when the z-scored signal exceeded 2 standard deviations for at least 15 ms and the animal’s speed was less than 4 cm/s. The detected SWR periods were then extended before and after the threshold crossings to include the time until the population trace returned to the mean value. Only SWRs with spikes from at least two tetrodes were included in replay analysis.

#### 4.1.3 Identifying SWRs during sleep

To identify SWRs in sleep states in the rest box, we followed the method described in Kay et al. [2016]. First, we extracted periods when the animals’ speed was < 4 cm/s, preceded by 60 s with no movement > 4 cm/s. Second, we detected REM periods following the method in Mizuseki et al. [2011]. Specifically, for all CA1 tetrodes, the ratio of Hilbert amplitudes (smoothed with a Gaussian kernel, *σ* = 1 s) of theta (5–11 Hz) to delta (1–4 Hz) filtered LFP was calculated, and then the mean was taken over tetrodes. REM periods were identified as sustained periods (10 s minimum duration) in which the theta to delta ratio was elevated above a manually set threshold (range: 1.2–1.8). Third, after excluding REM periods, LFPs from all available CA1, CA2, and CA3 tetrodes were squared, smoothed with a Gaussian kernel (*σ* = 0.3 s), and then z-scored and summed across all tetrodes. The sum trace was again z-scored to obtain an aggregate hippocampal LFP amplitude. A bimodality was observed in the histogram distribution of the aggregated LFP amplitude [Kay et al., 2016]. Fourth, SIA periods were defined as non-REM times in which the aggregate LFP amplitude was below the threshold separating the two modes on the histogram, and LIA periods otherwise. LIA periods were therefore high-amplitude LFP, corresponding to a hippocampal sleep state dominated by SWRs, frequently interrupted by periods of low-amplitude LFP, i.e., SIA. Only SWRs within those candidate sleep periods at least 90 s in duration, which contained extended (>5 s) continuous LIA periods, were included in sleep replay analysis.

### 4.2 State space decoder for replay and theta sequences

#### 4.2.1 The state space decoder

The state space model was described in detail in Denovellis et al. [2021]. One of the advantages of the state space decoder is its use of very small temporal time bins (2 ms) to perform a moment-by-moment estimation of the position representation (compared to the standard “Bayesian” decoder with >20 ms time bins). This property allows for the detection of rapid representational movement and therefore captures a range of movement dynamics, including super-diffusion (trajectories with interleaved slow progression and large jumps), stationary (unchanging representations), and diffusion (trajectories that progress at variable or constant speeds).

The state space model takes two inputs: the multiunit spikes described above and the linearized position of the animal. The linearization was done by converting the 2D position of the animal into a single line extending from the home well (central arm) to the edge of the left and right arms (central arm, right arm, and then left arm; see Fig.S1b). The linearized position was then binned into 3 cm spatial bins within each arm. Compared to decoding in a 2D scenario, such linearization significantly speeds up the decoding process. We adopted the “clusterless” version of the state-space decoder, which directly uses multiunit spikes and their spike waveform features to decode position without spike sorting. This clusterless decoder allows access to a larger pool of data than the spike-sorted decoder, providing more information about the extent of the event and greater certainty in the latent position representation[Denovellis et al., 2021].

We then ran the state space model independently for each session. Before decoding replay sequences and theta sequences, we first validated the model with the decoding accuracy of the animals’ locations during running (with animals’ speed > 4 cm/s). We found that the decoded position closely tracks the animal’s actual position (5-fold cross-validation; with the distance difference between the actual and decoded location having a median of 9.0 cm, and 95% CI of 8.2 − 9.8 cm), which is comparable to previous studies [Davidson et al., 2009, Farooq and Dragoi, 2019].

#### 4.2.2 Decoding of replay sequences

For the state space model of replay decoding, we built the encoding model using all multiunit spikes when the animal was moving > 4 cm/s and a 2-ms temporal bin. Three movement dynamics were included in the encoding model: 1) continuous movement dynamic—with an equal probability of moving one position bin forward or back; 2) stationary movement dynamic-staying in the same position; 3) fragment movement dynamic-with an equal probability of moving to any other possible position bin. The transition matrix, which defines how likely the movement state is to switch to another state versus persist in the same state, is set to have a 0.98 probability of persisting in the same state and 0.01 probability of switching to one of the other two states. After training the encoding model, we applied it to SWRs during both the awake immobile state and the subsequent sleep state. The main output of the model is the posterior probability of position, which is estimated by marginalizing the joint probability over the dynamics. This indicates the most probable “mental” positions of the animal based on the data. For replay diffusivity (Results. 2.4.1), we employed the method described in Methods. 4.9, by fitting the slope between replay step size and time duration on a log-log scale.

#### 4.2.3 Decoding of theta sequences

For the state space model of theta sequence decoding, we again built the encoding model using a 2-ms temporal bin. However, we only considered two movement dynamics: continuous and fragmented. The continuous dynamic was modeled by a random-walk transition matrix with a 6-cm standard deviation, and the fragmented dynamic was modeled by a uniform transition matrix. The probability of staying in either the continuous or the fragmented movement dynamic was set to 0.968, which corresponds to 62.5 ms of staying in the same movement dynamic on average, or roughly the duration of half a theta cycle. The probability of transitioning to the other movement dynamic is 0.032.

We used 5-fold cross-validation for decoding, in which we built the encoding model on 4 folds of the data and then decoded the sequences on the remaining fifth fold of the data. This ensures that the data used for constructing a given encoding model are not also used for decoding the representation. We repeated this for each fifth of the data. It is also noteworthy that we didn’t perform cross-validation for replay decoding since the training data and the testing data are naturally separated from each other. To calculate the average theta sequence length for each recording session (Results. 2.4.1), we averaged the theta sweep lengths (look-ahead distance plus look-back distance) across all LFP theta cycles during running.

### 4.3 The continuous attractor network (CAN) model with firing rate adaptation

The dynamics of the CAN model with firing rate adaptation is written as:

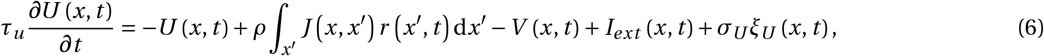

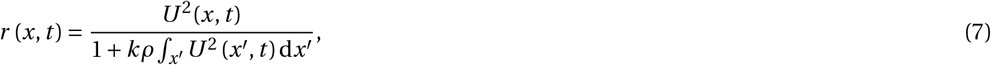

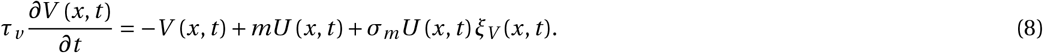

Eq. (6) describes the dynamics of presynaptic current of a neuron in the CAN with the firing rate (Eq. (7)) implemented as a global inhibition. Eq. (8) describes the dynamics of firing rate adaptation which is negatively fed back to the dynamics of the presynaptic current. *m* is the adaptation strength. *τ*_*u*_ is the neuronal time constant, and *τ*_*v*_ is the adaptation time constant which is much larger than *τ*_*v*_, highlighting the feature of slow feedback inhibition. *ρ* represents the place cell density covering the environment and *k* represents the strength of the global inhibition. *σ*_*U*_ and *σ*_*m*_ represent the noise amplitude applied to *U* and *V*, respectively.

The synaptic connection *J* (*x, x*_′_) is translation-invariant, which is written as:

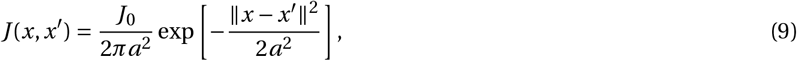

with *J*_0_ controlling the strength of the recurrent connection and *a* controlling the range of neuronal interaction. This translation-invariant form indicates that the synaptic strength of two place cells only depends on the distance between the locations of their place fields.

The location dependent sensory-motor input *I* ^*ext*^ (*x, t*) is modelled as a bump input conveying the information of the animal’s physical location to the CAN. It is expressed as:

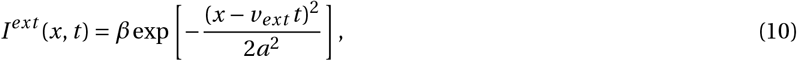

where *β* controls the input strength, and *v*_*ext*_ represents the moving speed of the artificial animal. For simplicity, we modeled the animal’s movement with a constant speed. To generate theta sequences in the model, we set *β* > 0 to mimic the interplay between external sensory input and the intrinsic dynamics from firing rate adaptation in the hippocampus (see Methods. 4.8 below). Conversely, to generate replay sequences, we set *β* = 0, simulating the absence of location-dependent sensory input during resting states, allowing the network state to evolve solely based on intrinsic dynamics.

Common parameters used to simulate replay-like sequences and theta sequences in the CAN model are summarized in Table. 1. For the key parameters, i.e., the adaptation strength *m* and the noise level *γ* (see Eq. (4)) for generating replay sequence with different diffusivity and theta sequences with different amplitudes are summarized in Table. 2. All simulations were conducted using the first-order Euler method.

**Table 1:** Common parameter settings of the 1D CAN model.

| Parameters | Values |
| --- | --- |
| **Common parameter setting** |  |
| Number of neurons: $N$ | 128 |
| Neuron density: $\rho$ | 20 |
| Recurrent connection range (Gaussian width): $a$ | 0.4 m |
| Synaptic connection strength: $J_0$ | 4 |
| Global inhibition strength: $k$ | 20 |
| Time constant of neural firing: $\tau_u$ | 1 ms |
| Time constant of firing rate adaptation: $\tau_v$ | 48 ms |
| Simulation time interval: $\delta t$ | 0.1 ms |
| **Common parameter setting for replay sequences** |  |
| Location dependent input strength: $\beta$ | 0 |
| Animals' running speed: $v_{ext}$ | 0 m/s |
| **Common parameter setting for theta sequences** |  |
| Location dependent input strength: $\beta$ | 0.01 |
| Animals' running speed: $v_{ext}$ | 1.5 m/s |

**Table 2:**
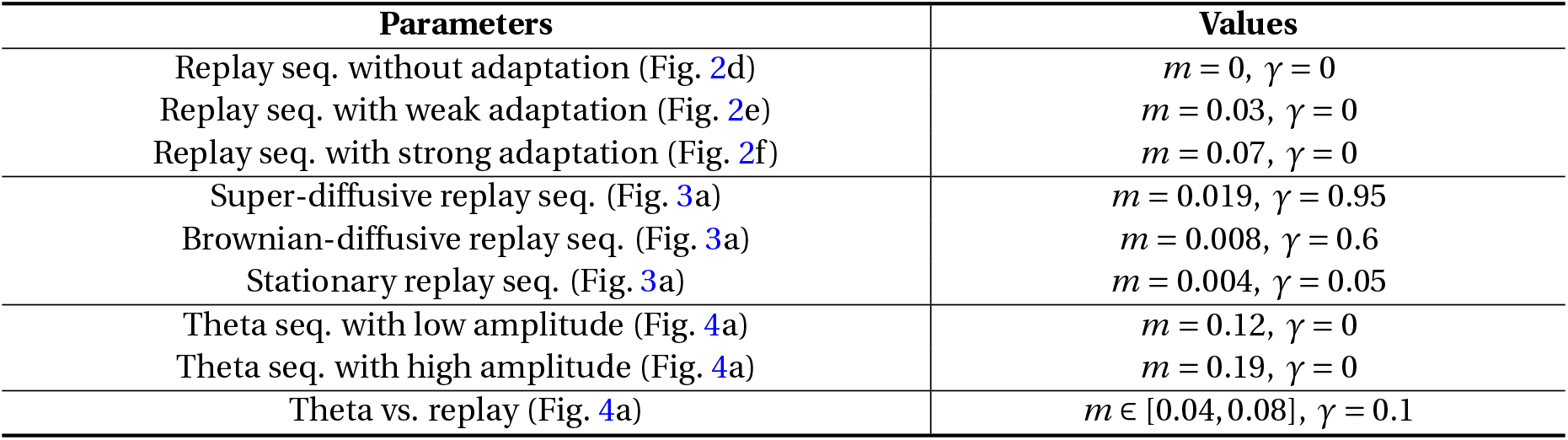
Parameter settings of the adaptation strength and the noise level in generating replay sequence with different diffusivity and theta sequences with different amplitudes.

### 4.4 Parameter conditions to generate bump activity in the CAN model

The global feedback inhibition (Eq.(7)) and the distance-dependent recurrent synaptic connection (Eq.(9)) prevent the neural activity from spreading in the CAN, and hence can result in a bump-like activity profile as the network state. However, from a mathematical perspective, this bump-like network state is not a trivial solution of the network dynamics. For instance, when the global inhibition is strong (large value of *k*), the activity bump may not survive. Following the theoretical analysis in Fung et al. [2008], we now derive the parameter conditions necessary to ensure the emergence of the bump state.

For simplicity, we consider that in the CAN model, there are only the recurrent connections and the global feedback inhibition, allowing us to investigate how these two factors affect the emergence of bump activity in the network. The network dynamics are then expressed as:

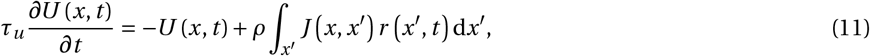

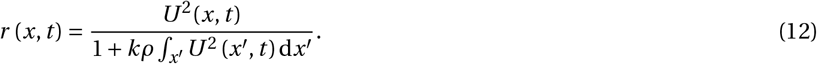

We assume the activity bump has the following profile (if it exists in the simple network):

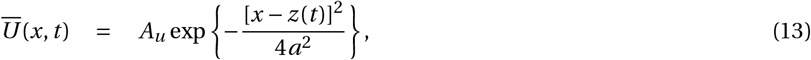

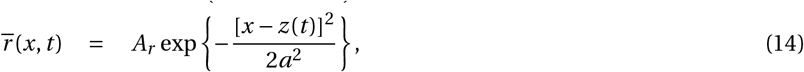

where *A*_*u*_, *A*_*r*_ represent the bump heights of *U* (*x, t*) and *r* (*x, t*), respectively. *z*(*t*) is the bump location, and *a* is the range of neuronal interaction. We then substitute Eqs. (13)&(14) into Eqs. (11)&(12) which gives:

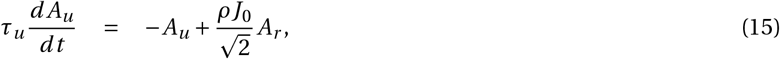

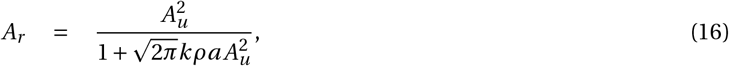

For the activity bump to exist in the CAN, the bump height should have a fixed positive value, which means *d A*_*u*_ /*dt* = 0, i.e., 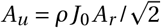 in Eq. (15). Combining it with Eq. (16), we obtain the solutions of *A*_*u*_ and *A*_*r*_, which are:

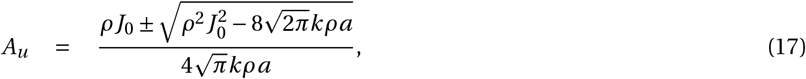

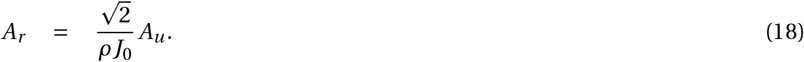

For *A*_*u*_ to exist, 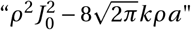 should be non-negative, which means 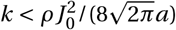 should be met. In summary, the condition that the CAN generates bump activity as its network state is that the global inhibition strength *k* is set smaller than a threshold determined by three other parameters in the CAN model, that is, the neuronal density *ρ*, the recurrent connection strength *J*_0_ and the neuronal interaction range *a*. In order to obtain a meaningful representation of the environment in the hippocampal place cell network (equivalent to localized bump activity in the CAN), we always choose *k* below the threshold throughout the paper.

### 4.5 Deriving the relationship between bump intrinsic speed and adaptation strength

Firing rate adaptation introduces instability to the activity bump, thereby causing intrinsic movement of the activity bump when there is no external input drive (Fig. 2c-f). A typical feature in Fig. 2c highlights that there exists a state transition boundary in the adaptation strength, below which the bump stays stationary, and above which the bump moves faster under stronger adaptation strength. We here theoretically analyze how the adaptation strength affects the movement of the activity bump.

To simplify the analysis, we consider a noise free network with *σ*_*U*_ and *σ*_*V*_ are all zero. We again assume the network activity has the following bump profile (now including the *V* (*x, t*)):

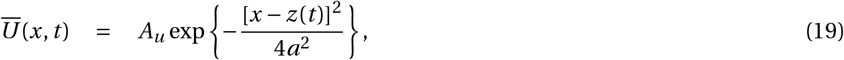

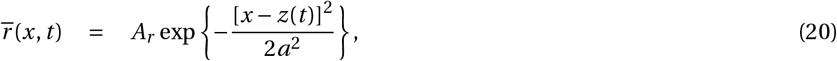

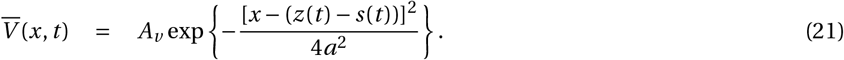

Here *A*_*v*_ is the bump height of the adaptation effect, and *z*(*t*) is the bump center. The intrinsic speed of the bump under firing rate adaptation is then described as *dz*(*t*)/*dt* (marked as *v*_*int*_ below). *s*(*t*) in Eq. (21) indicates the displacement between the position of the *U* bump and the *V* bump. Without loss of generality, we assume that the intrinsic movement of the bump is from left to right on the linear track, with 0 located at the left side. Therefore, *dz*(*t*)/*dt* > 0 always holds, indicating the bump travel to the right, and *s*(*t*) > 0 holds, indicating *V* (*x, t*) lags behind *U* (*x, t*) due to the slow dynamics in firing rate adaptation (*τ*_*v*_ ≫ *τ*_*u*_).

Following the analysis in Mi et al. [2014], we can solve the network dynamics by utilizing an important property of the CAN, that is, the dynamics of a CAN are dominated by a few motion modes corresponding to different distortions in the shape of a bump. Specifically, we can project the network dynamics onto these dominant modes and simplify the network dynamics significantly. The first two dominant motion modes used in the present study correspond to the distortions in the height and position of the Gaussian bump, which are given by

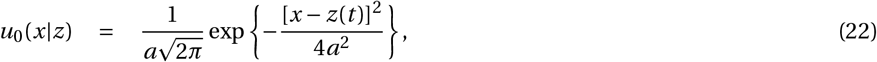

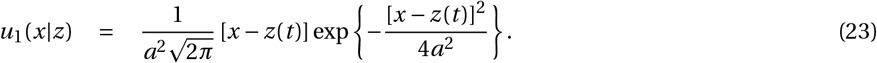

Projecting a function *f* (*x*) on a mode *u*(*x*) means computing _*x*_ *f* (*x*)*u*(*x*)*dx*. We first substitute the assumed network states (Eqs. (19)-(21)) into the network dynamics (Eqs. (6)-(8)) and then apply the projection method to simplify the dynamics, and then we obtain the intrinsic speed of the bump under firing rate adaptation *v*_*int*_, with

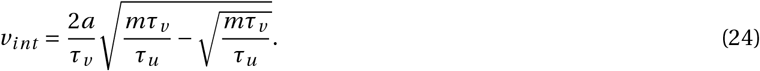

When *m* > *τ*_*u*_ /*τ*_*v*_, the intrinsic speed *v*_*int*_ is positive. We denote *τ*_*u*_ /*τ*_*v*_ ≡ *m*_0_ as the transition boundary, below which the activity bump stays stationary and above which, the activity bump moves intrinsically on the linear track. The intrinsic dynamics of the network depends on three factors in the network, i.e., the neuronal interaction range *a* (also controls the place field size), the time constant *τ*_*u*_ and *τ*_*v*_ and the adaptation strength *m*. For instance, when the adaptation strength increases, the bump moves faster (Fig. 2 c); when neurons have larger place field size, i.e., interacting more with each other, the bump also moves faster.

### 4.6 Deriving the position dynamics from the CAN model

We have shown that without noise, the activity bump exhibits intrinsic dynamics under the destabilization of firing rate adaptation. Now we investigate the bump dynamics under the joint effects of firing rate adaptation and network noise. The key idea behind the derivation is that the CAN dynamics (in the N dimensional neural space) can be simplified into the bump position dynamics and bump height dynamics over time (both of which are in one dimensional space) by a projection method [Fung et al., 2010] (similar to the analysis in Methods. 4.5 above). For the 1-dimensional dynamics, it is much easier to get the theoretical solution regarding how the position evolves over time, i.e., the quantification of replay diffusivity in the main text.

Specifically, we first substitute the assumed network states (Eqs. (19)-(21)) into the network dynamics (Eqs. (6)-(8)), and then project the network dynamics onto the bump height model (Eq. (22)). This operation gives us the dynamics of bump heights which are:

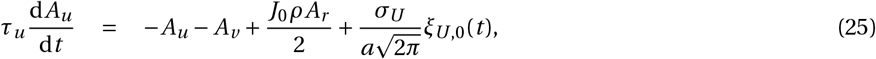

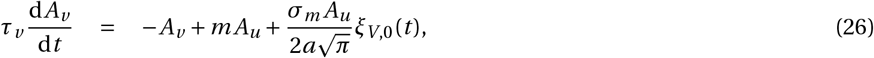

where *ξ*_*U*,0_(*t*) and *ξ*_*V*,0_(*t*) denote, respectively, the projected noises of *ξ*_*U*_ (*t*) and *ξ*_*V*_ (*t*) on the height mode, which are still Gaussian white noises of zero mean and unit variance.

Second, we project the network dynamics onto the position mode (Eq. (23)), and obtain the dynamics of bump positions which are:

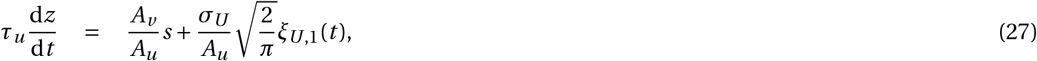

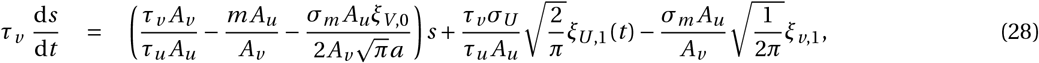

where *ξ*_*U*,1_(*t*) and *ξ*_*V*,1_(*t*) denote, respectively, the projected noises of *ξ*_*U*_ (*t*) and *ξ*_*V*_ (*t*) on the position mode, which are also Gaussian white noises of zero mean and unit variance.

Eqs. (25)& (26) can be described by the Fokker-Planck equations, which when solved, give the stationary distributions of *A*_*u*_ and *A*_*v*_ . Additionally, since *σ*_*U*_ and *σ*_*m*_ are relatively small, we ignore the variances of *A*_*u*_ and *A*_*v*_ and keep their mean values. Therefore, Eqs. (27)&(28) can be further simplified by replacing *A*_*u*_ and *A*_*v*_ with their mean values *Ã*_*u*_ and *Ã*_*v*_ .

Together with the approximation of *Ã*_*v*_ = *m *Ã**_*u*_ (according to Eq. (26)), Eqs. (27)&(28) can be written as:

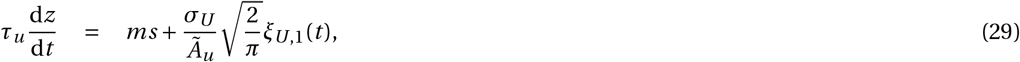

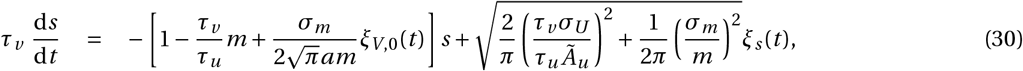

where *ξ*_*s*_ (*t*) is a Gaussian white noise of zero mean and unit variance (by combining the last two noise terms in Eq. (28)). We define 1−*τ*_*v*_ *m*/*τ*_*u*_ ≡ *µ*, which quantifies the normalized distance of the adaptation strength to the transition boundary *m*_0_, and define 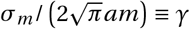, which quantifies the normalized noise amplitude. We also rewrite the noise terms in Eq. (29)& (30). After these operations, we obtain the position dynamics under the drive of both firing rate adaptation and noise fluctuations, which is:

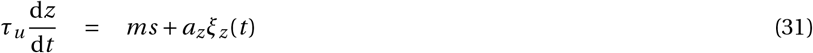

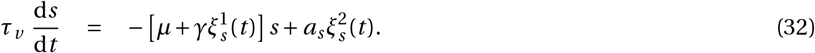

Eqs. (31)& (32) are typical Langevin dynamics, showing that the position dynamics *z*(*t*) is determined by a drift term reflecting the contribution of firing rate adaptation and a diffusion term reflecting the contribution of network noise. In fact, the position dynamics *z*(*t*) is a second order variable which depends on the dynamics of *s*, i.e., the displacement of bump *U* (*x, t*) and bump *V* (*x, t*). For instance, when the adaptation strength is set far below the transition boundary (small *m* and large *µ*), *s* decays quickly to zero, and *z* is determined only by the noise diffusion term *a*_*z*_*ξ*_*z*_ (*t*), and hence exhibit the dynamics of Brownian motion; when the adaptation strength is set near the transition boundary (large *m* and small *µ*), *z* is determined by both the drift and the noise diffusion term, and hence exhibits super-diffusive dynamics.

It is noteworthy that the noise term 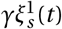 in Eq. (32) is necessary for generating the super-diffusive dynamics. If *γ* = 0, Eq. (32) becomes an Ornstein–Uhlenbeck (OU) process, and the stationary distribution of *s*(*t*) has a Gaussian form, which leads to two additive noises in Eq. (31), and the position dynamics only exhibits Brownian motion. We will quantify the diffusivity in a power law expression of the step size below.

### 4.7 Obtaining the probability distribution of the step size from the position dynamics

The position dynamics (Eq. (31)& (32)) are a second order process. Therefore, to solve the position dynamics of *z*(*t*), we first solve the dynamics of *s*(*t*). Following the analysis in Dong et al. [2021], we can describe the dynamics of *s*(*t*) as a Fokker-Planck equation and obtain the probability distribution of *s*(*t*) which has the form of a power law:

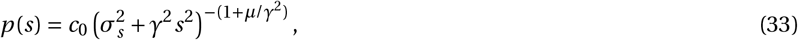

where 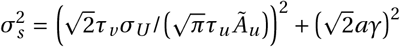 and *c*_0_ is a normalization constant. The dynamics of *z*(*t*) (Eq. (31)) shows that the step size of the activity bump in Δ*t* is:

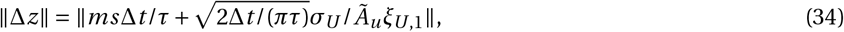

with Δ*t* → 0. Therefore, by replacing *s* with its stationary distribution given by Eq. (33), we obtain the power law distribution of the step size ∥Δ*z*∥, which is written as,

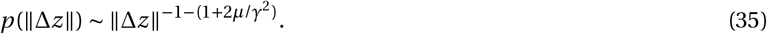

Eq. (35) shows that increasing the adaptation strength (decreasing the value of *µ*) and/or increasing the noise level (increasing the value of *γ*) can increase the probability of large step size, that is, the probability of long-jump movements of the activity bump on the linear track.

### 4.8 Generating theta sweeps in the CAN model

Theta sequences have been hypothesized to result from the interaction of external location dependent sensory input and the intrinsic network dynamics [Drieu and Zugaro, 2019, Tsodyks et al., 1996]. Therefore, following previous work [Chu et al., 2024], we generated theta sequences in the CAN model by activating the location-dependent sensory input *I*_*ext*_ (*x, t*) (Eq. (10)). The external input is modeled as an activity bump traveling with a constant speed, simulating the update of the animal’s physical location in the environment. The interaction of external input and the intrinsic dynamics creates a push-pull effect on the activity bump: as the animal advances, the external input exerts a constant pull effect on the activity bump, attracting it back to the current physical location, while slow feedback inhibition (adaptation) pushes it away from the current physical location. This results in theta-like sweeps of the location representation as the animal explores the environment (see Fig. 4a and Table. 1 for parameter settings). Intriguingly, akin to how the adaptation strength governs replay diffusivity in the CAN model, it also regulates the sweep amplitude of generated theta sequences. Specifically, stronger adaptation results in a larger sweep amplitude, as illustrated by comparing Fig. 4a left and right. This phenomenon arises from the increased intrinsic mobility in the activity bump associated with stronger adaptation, causing it to sweep further during the push effect.

### 4.9 Diffusion exponent analysis of replay trajectories

The power law distribution of step size *p*(∥Δ*z*∥) ∼ ∥Δ*z*∥_−1−*α*_ (Eq. (3)) rigorously characterizes the diffusivity of a movement trajectory. However, the condition of obtaining the probability distribution is as Δ*t* → 0. This implies that, to quantify the diffusivity of replay trajectories, we need to fit a power law distribution of the step size in infinitesimal time bins. However, in experimental data analysis, the decoding time bin is typically set between 2-20 ms (depending on the decoding algorithm), posing a challenge for the fitting process. Therefore, we followed previous work [Stella et al., 2019, Krause and Drugowitsch, 2022] to quantify the diffusivity of a movement trajectory as the relationship between distance and time, which is expressed as:

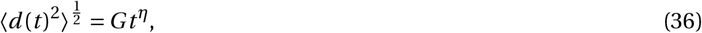

where *d* is the distance between two points in time within a replay trajectory, *t* is the time elapsed between those two time points, *G* is a constant describing the scale of diffusion, and *η* is the diffusion exponent. To quantify the diffusivity of replay trajectories in experimental data, we plot the relationship between *d* (*t*) and *t* in log-log space, and use linear regression applied to this log-log plot to find the slope, which corresponds to the diffusion exponent *η*. A slope (or *η*) of 0.5 corresponds to Brownian diffusion, whereas a slope (or *η*) greater than 0.5 corresponds to super-diffusion.

Instead of quantifying the diffusivity of each replay trajectory (which contains only a few time bins, and therefore does not have enough data points for fitting), we quantify the overall diffusivity of replay trajectories in one recording session by merging all step sizes and their corresponding time bins from all replay trajectories in the recording epoch, and calculate one diffusion exponent by linearly fitting these two variables in the log scale. This allows us to quantify the overall diffusivity of replay dynamics from one recording epoch. Specifically, we calculate the distance *d*_*k*_ (Δ*t*_*j*_) between all pairs of decoded positions separated by all multiples of the time bin used for trajectory decoding Δ*t*_*j*_ /*δt* where Δ*t*_*j*_ is the time elapsed between decoded positions and *δt* is the unit decoding time bin. *k* is the *k*_*th*_ replay trajectory. This resulted in a set of distance-time pairs in one recording session:

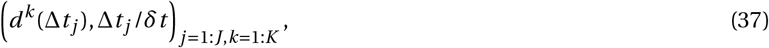

with *J* the number of elapsed time bins and *K* the number of replay trajectories in one recording session. We then calculate the mean distance 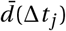 for each multiple *j* of the time bin by taking the average over all trajectories and fit a linear regression model to 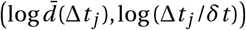, which gives the estimate of the diffusion exponent *η* for the recording session as the slope of the regression.

The two descriptions of diffusivity (Eq. (3) and Eq. (36) are interrelated to each other. Movement trajectories with the diffusion exponent *η* = 0.5 or the power law exponent *α* ≥ 2 follow Brownian diffusion, while movement trajectories with the diffusion exponent *η* > 0.5 or the power law exponent 1 < *α* < 2 follow super-diffusion. A slight difference is that Eq. (3) with 1 < *α* < 2 describes a sub-type of super-diffusion, that is, Lévy flights (see our previous work [Dong et al., 2021]), which is composed of frequent local motion and intermittent long-jump motion; while Eq. (36) with *η* > 0.5 describes a more general super-diffusion, which includes not only Lévy flights, but also replay dynamics with a constant fast speed. Since both cases have been observed in previous works [Pfeiffer and Foster, 2015, Krause and Drugowitsch, 2022, Denovellis et al., 2021], we use the diffusion exponent (Eq. (36)) to describe experimental results while use the power exponent (Eq. (3)) to describe the model results.

### 4.10 Measuring the place field size index (population vector correlation analysis)

To check the confounding factor of place field size in contributing to the positive correlation of replay diffusivity and theta sweep length, we calculate the place field size index for each recording session via the population vector correlation analysis (PVC) [Battaglia et al., 2004]. The PVC is a measure of how quickly the spatial firing patterns change with distance (i.e., how quickly the vectors decorrelate), and therefore implies the average size of place fields. Since we used the clusterless version of the state space model, we treat each recording unit as a “cell”. For each recording session, the rate matrix was constructed by arranging all the recording units into a two dimensional matrix 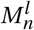, where the unit identities (*n* in total) are represented on the first dimension and the spatial intensity of unit activity (*l* bins in total) on the second dimension. As in building the encoding model for the state space decoder, the spatial intensity is calculated using the data when the animal’s speed is more than 4 cm/s. To provide an estimate of the similarity of the hippocampal neuronal ensemble code, we calculate the Pearson correlation coefficient for each pair of *M* -dimension population vectors at two spatial bins, and obtain a *l* ×*l* correlation coefficient matrix (Fig. S7b). Since the animals perform a spatial alternation task between the left and right arm, we further split the correlation coefficient matrix into two matrices, one represents the spatial similarity from the central to the left arm and the other represents the spatial similarity from the central to the right arm (Fig. S7b). The mean similarity value along the diagonal is calculated and further smoothed with a Gaussian kernel with *σ* = 5 for both arms, and then averaged across two arms. This results in a population vector correlation curve as a function of the diagonal offset (Fig. S7c). Finally, the place field index is calculated as the slope from the peak to the trough value on the curve, which measures how quickly the PVC decays among the spatial bins. A large place field index (large slope) represents a quick decay of the PVC value along the spatial bins, and hence indicates smaller place fields in the current recording session.

### 4.11 CAN model with external slow-gamma oscillation

To show the phase-locking phenomenon of movement step sizes and neural activity to the slow-gamma oscillation during sleep SWRs, we consider a CAN with an external input oscillating at the slow-gamma rhythm, which is written as:

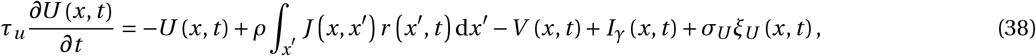

where *I*_*γ*_ (*x, t*) has a sinusoidal waveform given by:

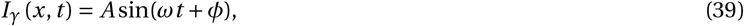

with *A* representing the amplitude, *ω* the angular frequency and *φ* the initial phase of the slow-gamma rhythm. Without loss of generality, we simply set *A* = 0.5, *ω* = 30, and *φ* = 0 during the simulation. For other parameters, see Table. 1.

### 4.12 Spectral analysis of the CAN sequence generator

#### 4.12.1 Convert the CAN dynamics into a sequence generator

In the CAN, the activity bump vector *U* containing a finite population of N neurons evolves according to Eq. (6)-(8), and can be interpreted as the distribution of current state estimate under idealistic setup (e.g., given spikes simulated from the network dynamics, Eq. 2). Hence the CAN can be conceptualized as a continuous-time Markov process (e.g., a generator model) with specific constraints so that the state vector represents a valid probability distribution which evolves over time (Fig. 3c) [McNamee et al., 2021]. This analogy allows us to perform spectral analysis of the evolution matrix (i.e., the generator) and derive the condition for generating distinct sequential dynamics.

To derive the generator model, we first vectorize the network dynamics in Eq. (6)-(8) as follows:

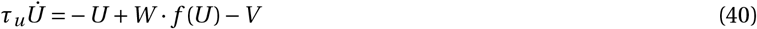

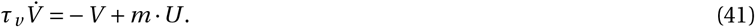

Here, *U*, *r, V* are now population vectors with each dimension representing the states of a place cell. Since the global inhibition (Eq. (7)) restrict the activity bump from spreading without changing the location of the bump, *f* (*U*) can be treated as an identical mapping as *f* (*U*) = *IU* . Furthermore, since the adaptation bump V lags behind the activity bump U with a displacement *s* (see Eq. (32) and Fig. 3d) but shares the same Gaussian profile as *U*, it can be treated as a shift version of *U* with *V* = *CU*, where *C* is a sub-diagonal matrix with zeros except the *s*-th upper diagonal with value of *ϵ*. Hence, from a normative account, we can interpret *V* as an exponential moving average of past state distributions for constraining smoothness of the dynamics of *U*, which lead to constant sub-diagonal perturbation to the canonical evolution matrix (without feedback inhibition). The detailed value of *ϵ* depends on the adaptation strength *m* and the time scale *τ*_*v*_, with larger adaptation strength/smaller time scale leading to larger values of *ϵ*. But for simplicity, we consider a fixed value of *ϵ* below. With these assumptions, the evolution of the states can be simplified as:

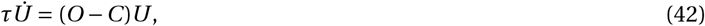

with *O* = *W* − *I* . We will focus our analysis on the 1D ring track environment for simplicity, where the transition matrix *O* is a circulant matrix.

#### 4.12.2 Spectral analysis of the perturbed transition matrix

For notational simplicity, we define *O*^*s*,*ϵ*^ = *O* −*C*^*s*,*ϵ*^ as the perturbed transition operator, where as above, *C*_*s*,*ϵ*_ represents the sub-diagonal matrix with constant *s*-th upper diagonal (of value *ϵ*) and zero everywhere else. Our key observation is that both the original and perturbed transition matrices are circulant matrices, hence sharing the same set of eigenvectors, with perturbed eigenvalues [Bracewell, 1986, Yu et al., 2020]. Specifically, the (normalised) eigenvectors for the circulant transition matrix over a circular state space of *N* states are the set of Fourier modes (Fig. 3e).

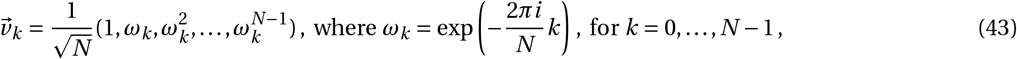

And the corresponding eigenvalues are the discrete Fourier transforms for the first row of *O*.

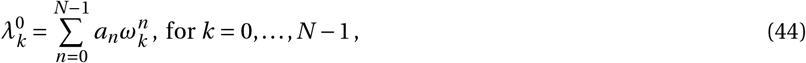

where 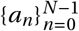 is the set of elements that uniquely define the circulant matrix (first row or column of the matrix).

Following the above Fourier analysis, we can analytically compute the perturbation to individual eigenvalues following the (*s, ϵ*)-perturbation to the transition operator.

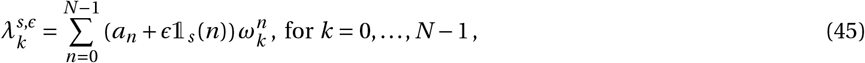

where 𝟙_*s*_ (*n*) is the indicator function that equals to 1 when *n* = *s* and 0 otherwise. Hence the (*s, ϵ*)-perturbation to the *k*-th eigenvalue is as following.

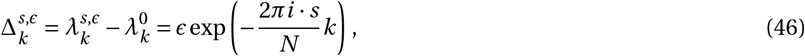

As a reminder, the eigenvectors remain unchanged. Hence, the perturbation effects on the transition dynamics are fully characterised by perturbations in eigenvalues (Fig. 3f and Fig. S2). Increase / decrease in (absolute values of) eigenvalues lead to amplifying / damping effects in the spatial distribution modelled by the corresponding eigenvectors. Hence, increase in eigenvalues corresponding to eigenvectors of larger spatial scales lead to increased tail probabilities in the transition dynamics, leading to higher probability for sampling distant locations (super-diffusivity). The precise effect can be qualitatively illustrated via examining the ratio between absolute values of perturbed and original eigenvalues (Fig. S2a&b). Depending on the offset associated with the perturbation, there exists a spectrum of oscillatory patterns dependent on the perturbation offset. Here we take two extreme cases for demonstration and perform full analysis of a simplified case – “Lazy” random walk below. With small offset (*s* = 5), eigenvalues of larger spatial scales (corresponding to lower-frequency Fourier modes) are dampened, and the ratios in absolute value with respect to original eigenvalues increase with the decrease in spatial scales, leading to decrease in tail probabilities, hence local diffusion is more likely. On the other hand, with larger offset, increase in eigenvalues of larger spatial scales precedes decrease in eigenvalues of smaller spatial scales, amplifying transition probabilities associated with the distant states, hence it becomes more likely for super-diffusive behaviour to emerge. Intermediately small perturbation offsets (e.g., *s* = 15, Fig. S2b) lead to reversed oscillatory pattern comparing to that of *s* = 50. In addition, we could analytically derive the precise boundary values for the offset at which the animal switches from local-to super-diffusion (see below), hence providing the ability to precisely track the nature and dynamics of replay sequences.

#### 4.12.3 A simple case of “Lazy”-random walk for further analysis of the eigenvalue rescaling problem

For the simplicity of quantitative analysis, we consider a “Lazy”-random walk transition matrix (for analytical computation of the perturbed eigenvalues, Fig. S2c).

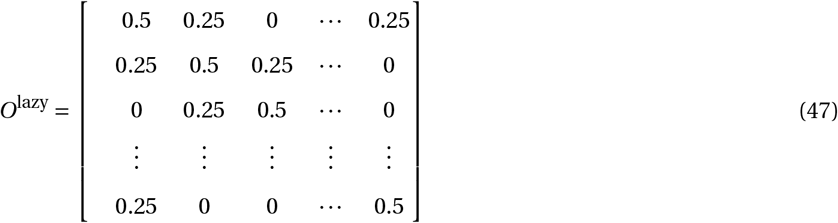

We could analytically compute the eigenvalues of *O*^lazy^.

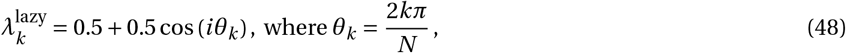

Perturbations to the principal eigenvector (second principal due to the constant Fourier mode corresponding to *k* = 0) has the most dominant effect on the resulting dynamics, we hence focus our analysis on 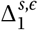 for all *s*.

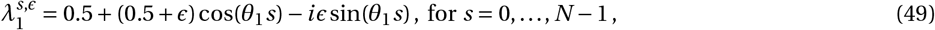

Examining the set of perturbed eigenvalues corresponding to different offsets reveals that the transition from dampened to amplified principal eigenvalue happens as *s* = 25 (and *s* = 75 given the symmetry, Fig. S2d). We additionally verify this under the Gaussian diffusion policy studied in the main text, which conforms well with our theoretical analysis (Fig. S2d). Note that the precise boundary is dependent on a number of factors, including the value of *ϵ*, which we have assumed to be constant in the current instance.

As discussed in Results 2.2, varying the perturbation offset leads to a spectrum of oscillatory patterns on the ratio between perturbed and original eigenvalues. To maximally demonstrate our intuition, we only studied two extreme cases in the main text. However, the complete range of oscillatory patterns are much more heterogeneous and harder to analyse (Fig. S2e), involving complicated interplay amongst *s, ϵ*, and spatial scale of eigenvalues (*k*, Equation 44). The main intuition is that for intermediately small offsets (up to symmetry), the oscillatory patterns are reversed comparing to larger offsets (Fig. S2b&e), i.e., decrease in eigenvalues of larger spatial scales lead increase in eigenvalues of smaller spatial scales, hence the resulting dynamics will more likely be local diffusion. We leave more concrete analysis of such spectral behaviour for future studies.

Note that for simplicity, we assume symmetric transition matrix (for both Gaussian diffusion and lazy random walk), but note that our analysis generalises to arbitrary asymmetric and circulant transition matrices (e.g., corresponding to non-trivial translation) so long as the circulant symmetry is preserved given the perturbation.

## Author Contributions

ZJ, TC, XD, NB and SW conceptualized and designed the research. ZJ analyzed the experimental data with the input from NB. ZJ, TC, XD, CY performed theoretical analysis and simulations. ZJ, TC, XD, DB, NB and SW interpreted the results. NB supervised the analysis of experimental data and SW supervised the analysis of theoretical modeling. ZJ, NB and SW wrote the manuscript with the input from all the other authors.

## Acknowledgements

We thank Eric Denovellis for sharing the code of the state space decoder. We thank Loren Frank and colleagues for making the experimental data online. We thank Kenneth Kay, Thomas Wills, Mattias Horan, Wentao Qiu, Wenhao Zhang for valuable discussions. This work was supported by: a Science and Technology Innovation 2030-Brain Science and Braininspired Intelligence Project (STI2030-Major Projects 2021ZD0200204, SW), a Wellcome Principal Research Fellowship (NB), a UKRI Frontier Research Grant (EP/X023060/1, DB) and an International Postdoctoral Exchange Fellowship Program (PC2021005, ZJ).

## Code and data availability

Code for reproducing all the results in the main text will be available before publication. All experimental data are taken from the Collaborative Research in Computational Neuroscience (CRCNS) hc-6 dataset contributed by Loren Frank and colleagues [Karlsson et al., 2015]. They are publicly available at: https://crcns.org/data-sets/hc/hc-6

## Supplementary figures

**Figure S1:**
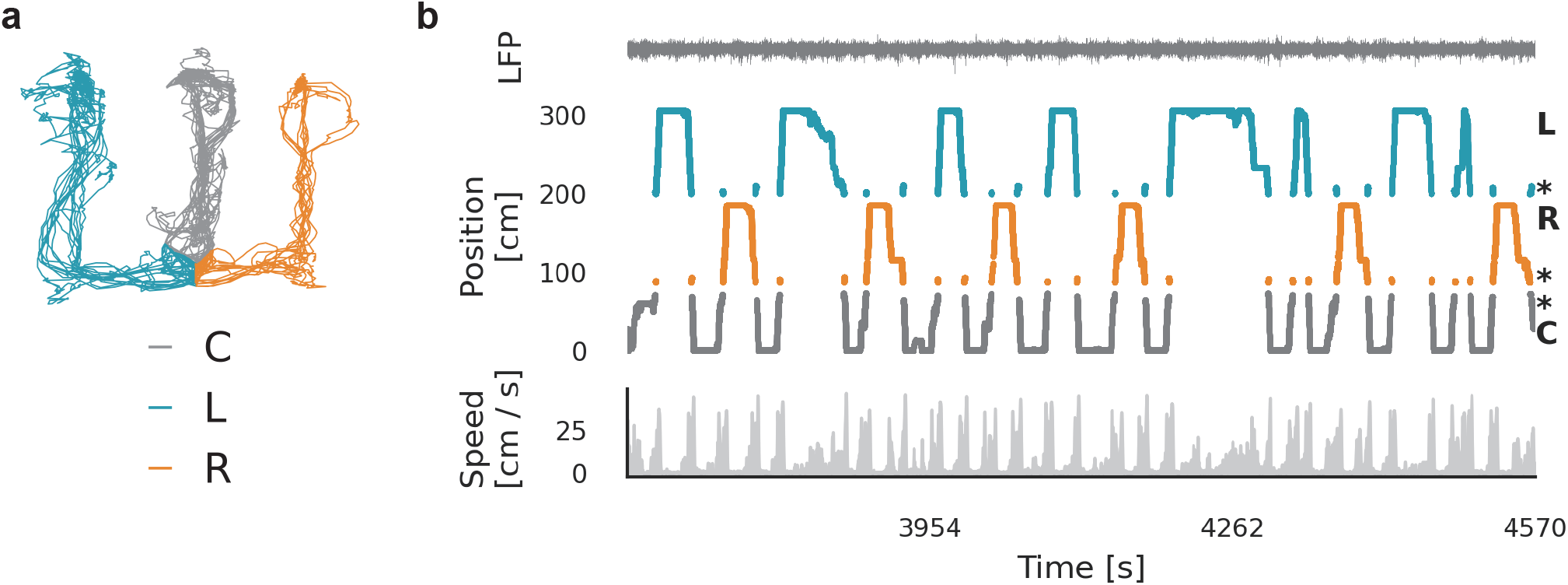
The W-shape spatial alternation task from [Karlsson et al., 2015]. **(a)**, the running trajectories from an animal during a recording session. Trajectories on the central arm, left arm and right arm are colored in grey, blue and orange, respectively. **(b)**, top panel: local field potential from a CA3 tetrode. Middle panel: the animal’s trajectories alternating on the central-left-central-right-central arms over time, with the colors same as in **(a)**. Bottom panel: the animal’s running speed over time.

**Figure S2:**
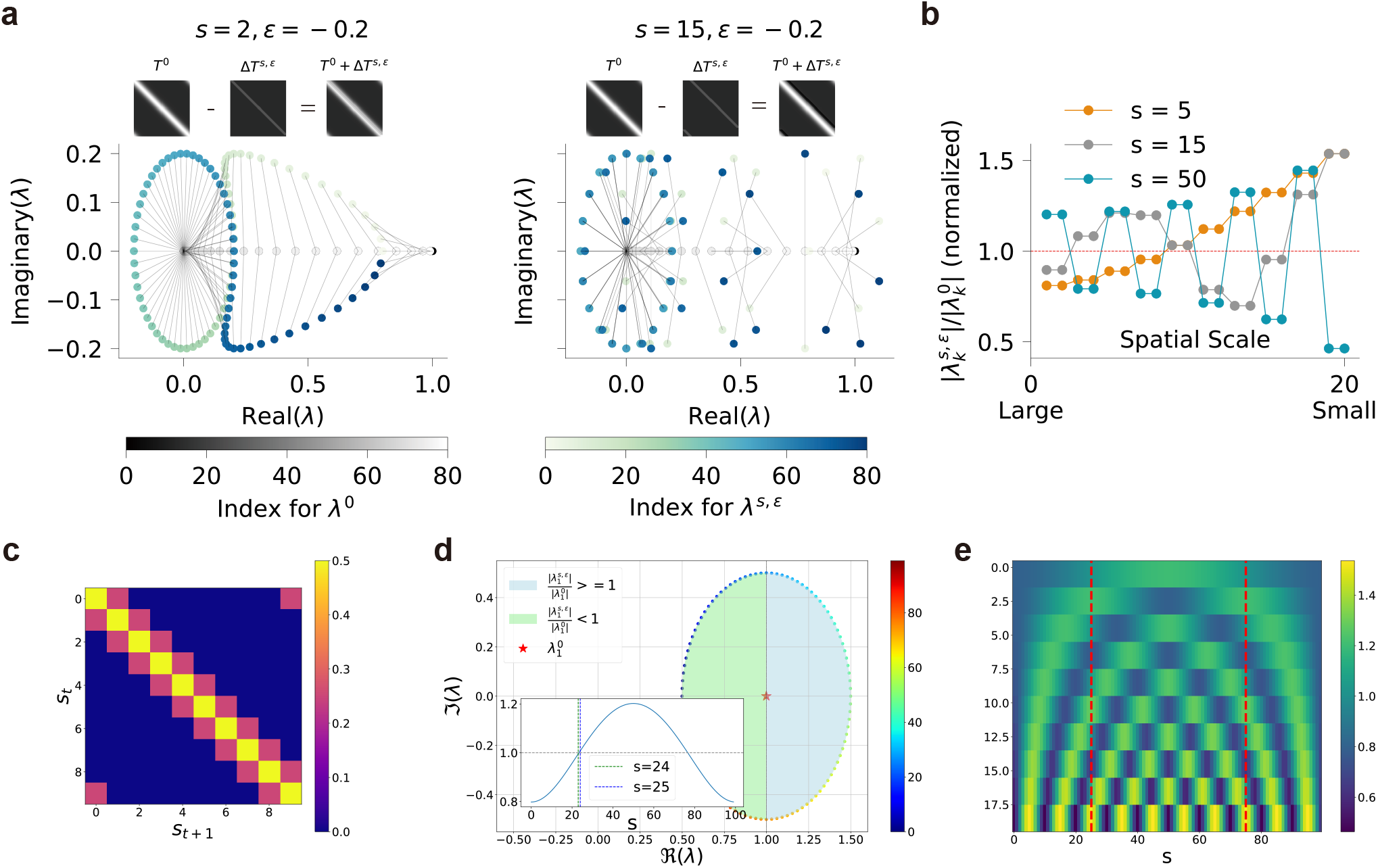
The whole spectrum of the perturbed transition matrix. **(a)**, Qualitative demonstration of perturbation to the eigenvalues following sub-diagonal perturbation to the transition dynamics. The perturbation patterns are heterogeneous with respect to perturbation offset, examples corresponding to *s* = 2 (left) and *s* = 15 (right) show vastly different distribution (symbols). grey bars link the perturbed eigenvalues with the original eigenvalues (without perturbation, lying on the real axis). **(b)**, rescaled eigenvalues (normalized by original eigenvalues) after the perturbation at different offsets (s=5: orange; s=15: grey and s=50: blue; top 20 eigenvalues are shown for better illustration). Theoretical analysis of oscillatory patterns in eigenvalue perturbations as a function of perturbation offsets. **(c)**, the “Lazy”-random walk transition matrix for analysis simplicity with 25% probability of transitioning to the left or the right location. **(d)**, perturbation of the second eigenvalue with different offset values under the simple “Lazy”-random walk scenario. 25 < *s* < 75 gives amplified values of the eigenvalue (blue area) and *s* < 25 and *s* > 75 gives dampened values of the eigenvalue (green area). **(e)**, the complete spectrum of the eigenvalue rescaling under the simple “Lazy”-random walk scenario. x-axis represents the perturbation offset, each row is one rescaled eigenvalue (> 0 means amplifying; < 0 means dampening). Re-scaling of the first 20 eigenvalues are shown for demonstration. Red dashed lines mark the boundary of amplifying/dampening effect for the eigenvalue showed in **(d)**.

**Figure S3:**
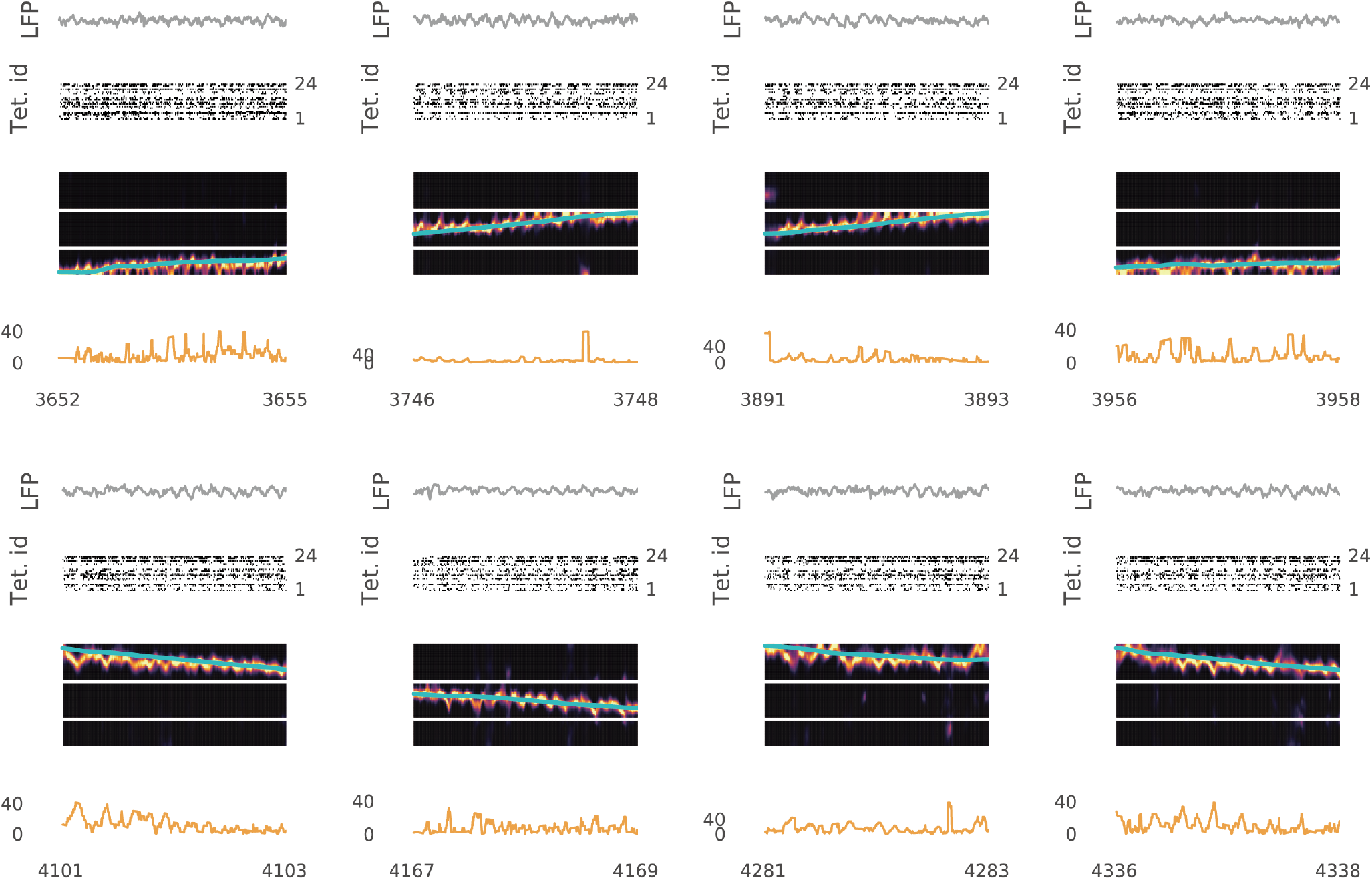
More examples of decoding theta sequences during running. From top to bottom in each example: the LFP signal from a CA1 tetrode; the multiunit activity; the posterior probability map; the offset distance between the decoded position and the actual position as a function of time (in seconds).

**Figure S4:**
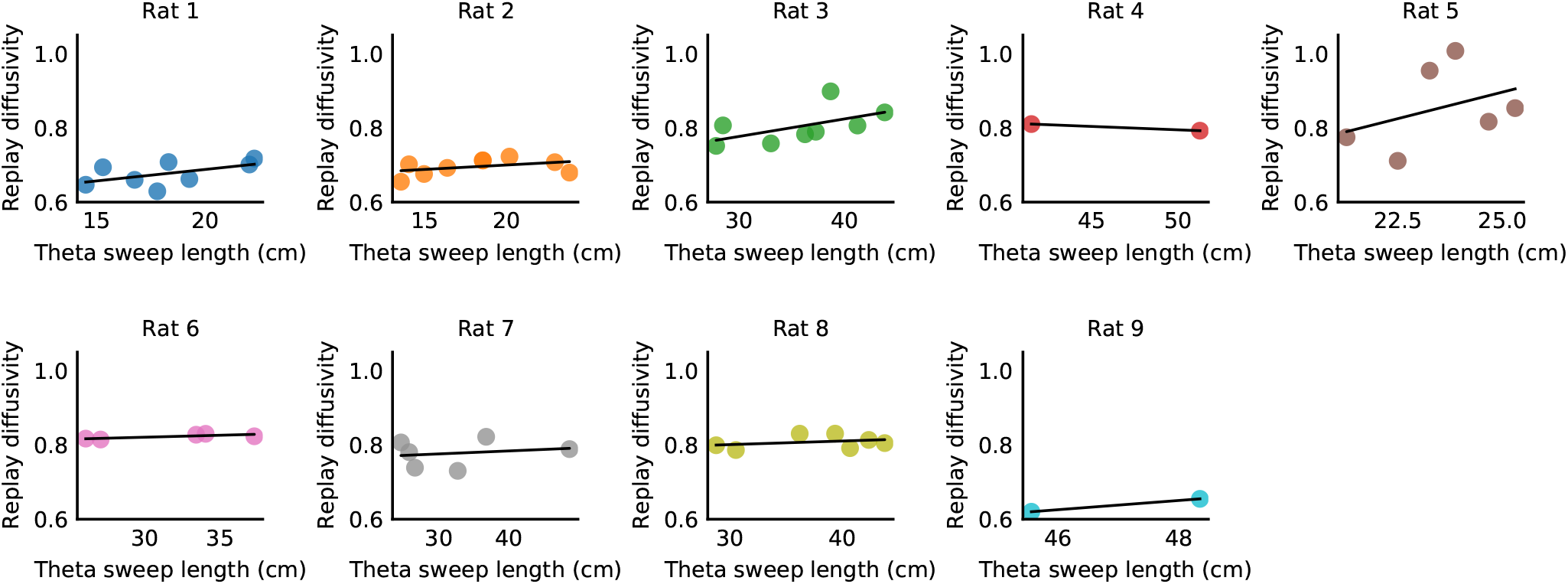
Correlation between replay diffusivity and theta sequence length across individual animals (9 in total). Each dot represents the diffusion exponent and the average length of theta sequences from a single recording day per animal. Note that while no significant correlation is observed, likely due to limited data points, there is a discernible trend suggesting a positive correlation.

**Figure S5:**
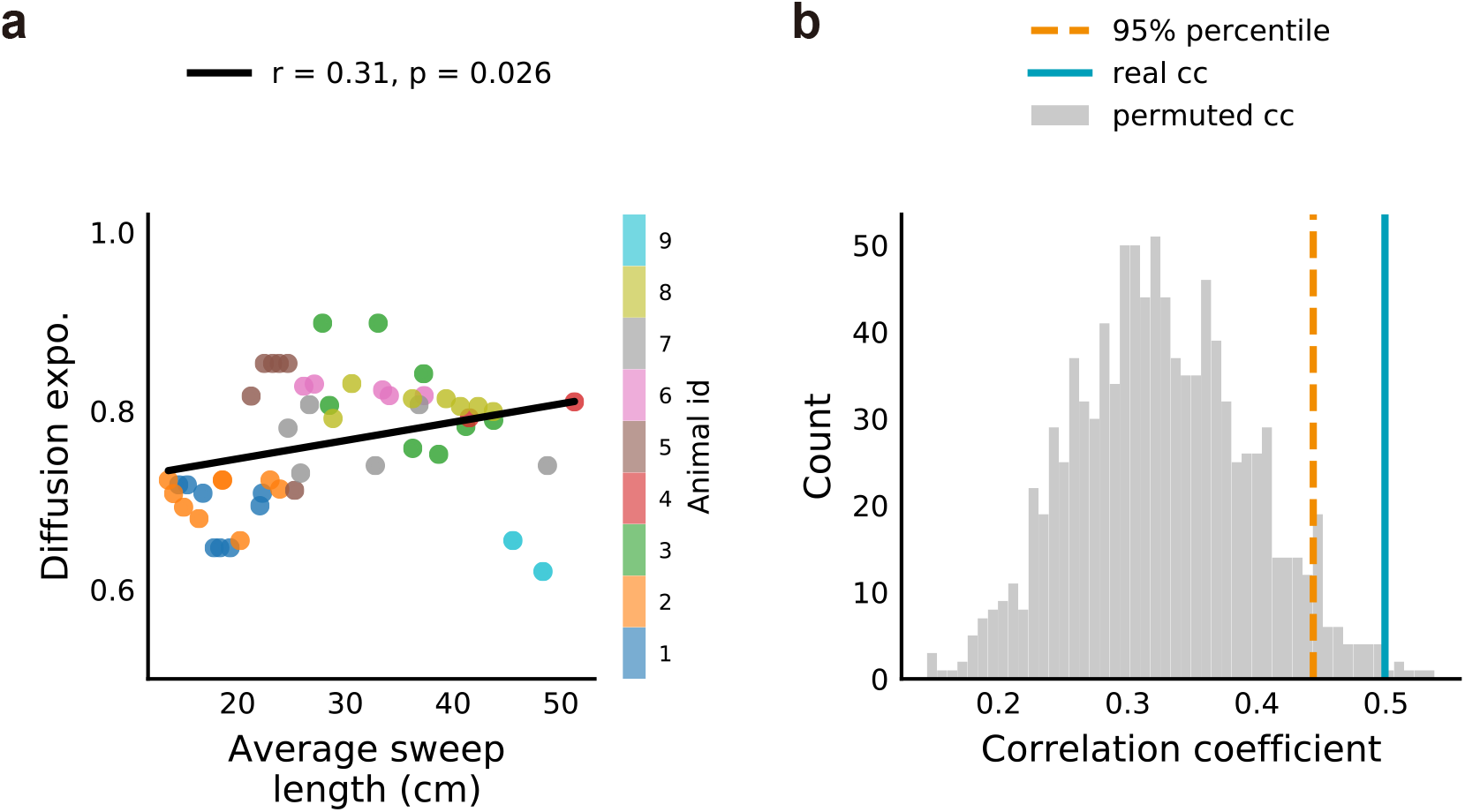
Shuffling the replay diffusivity across recording days within the same animal. **(a)**, an example of the correlation between the theta sweep length and the shuffled replay diffusivity (by randomly sampling an replay diffusivity from another recording day of the same animal). **(b)**, the histogram of the correlation coefficients between shuffled replay diffusivity and theta sweep length (1000 shuffles). The real correlation coefficient between replay diffusivity and theta sweep length is marked as the blue vertical line, and the 95% percentile of the shuffled correlation coefficients is marked as the orange dashed line.

**Figure S6:**
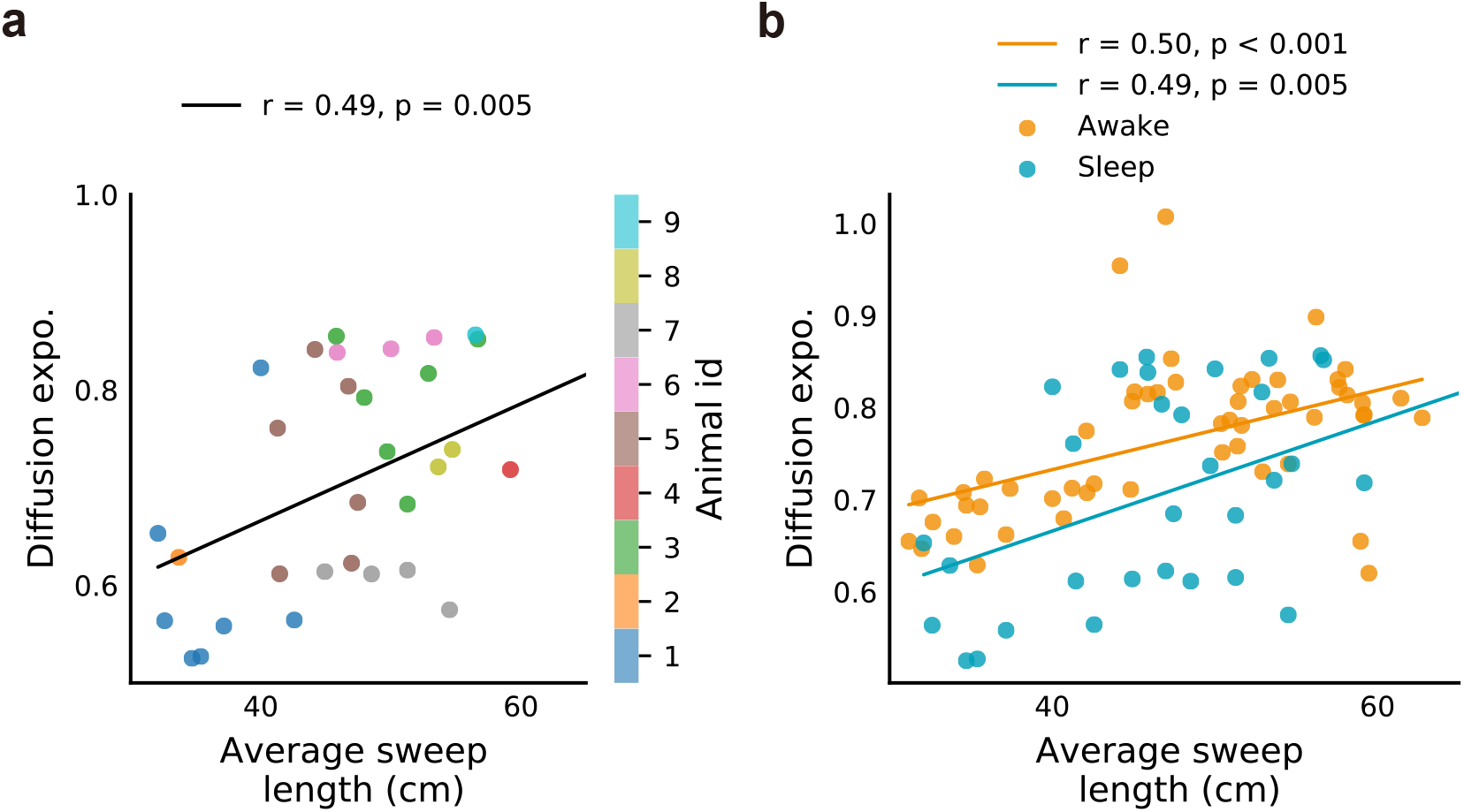
Correlation between theta sweep length and the diffusion exponent during subsequent sleep replay. **(a)**, each point indicates the averaged length of theta sequences and the diffusion exponent during subsequent sleep replay from one recording session, with different colors representing data from different animals. **(b)**, the correlation between theta sweep length and the diffusion exponent of awake replay in the same recording session (orange), and the correlation between theta sweep length and the diffusion exponent of subsequent sleep replay (blue). Note that the correlation coefficients are similar and the diffusion exponent in awake replay is generally higher than that during subsequent sleep replay.

**Figure S7:**
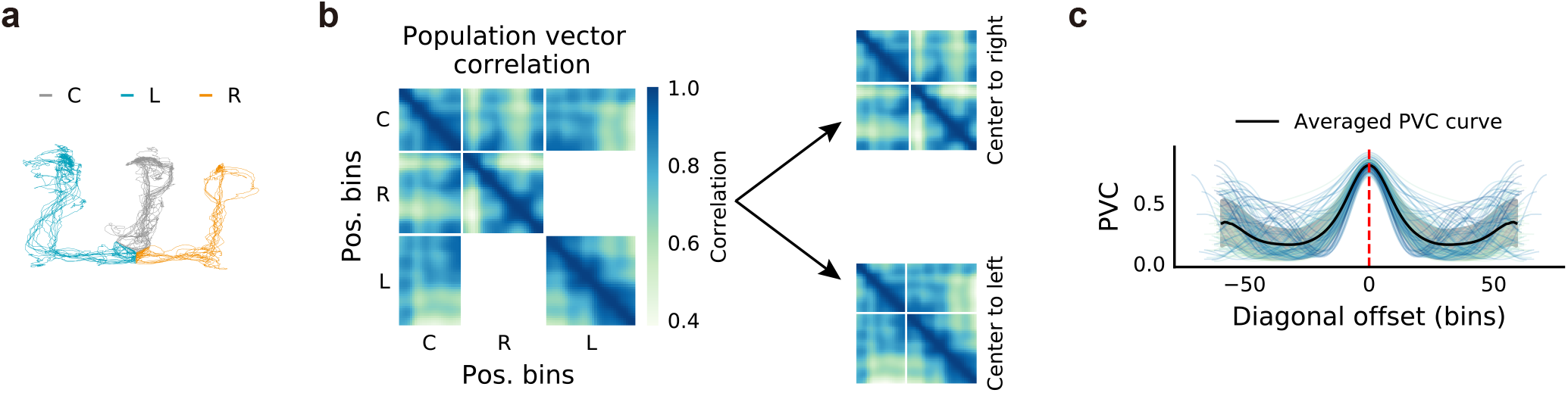
Measuring the place field size index (population vector correlation analysis). **(a)**, the movement trajectory of an animal in one running session, with three arms coded by different colors. **(b)**, the population vector correlation matrix shown in linearized spatial bins of center-right-left arms (left) and in separated center-right arm (top right) and center-left arm (bottom right). **(c)**, the population vector correlation curve as a function of the diagonal offset of the correlation matrix. Each curve represents the correlation along the diagonal of one recording session, and the dark line marks the averaged curve. The correlation value is highest when correlation the population firing vector in the same spatial bin (auto-correlation), and slowly decays as the two spatial bins become further apart to each other (larger offset). Toward the end of the offset, there is an increase in correlation values, which may reflect an over-representation of reward locations at the end of the three arms or the geometrical similarity of the end of the three arms in the 2D space.

**Figure S8:**
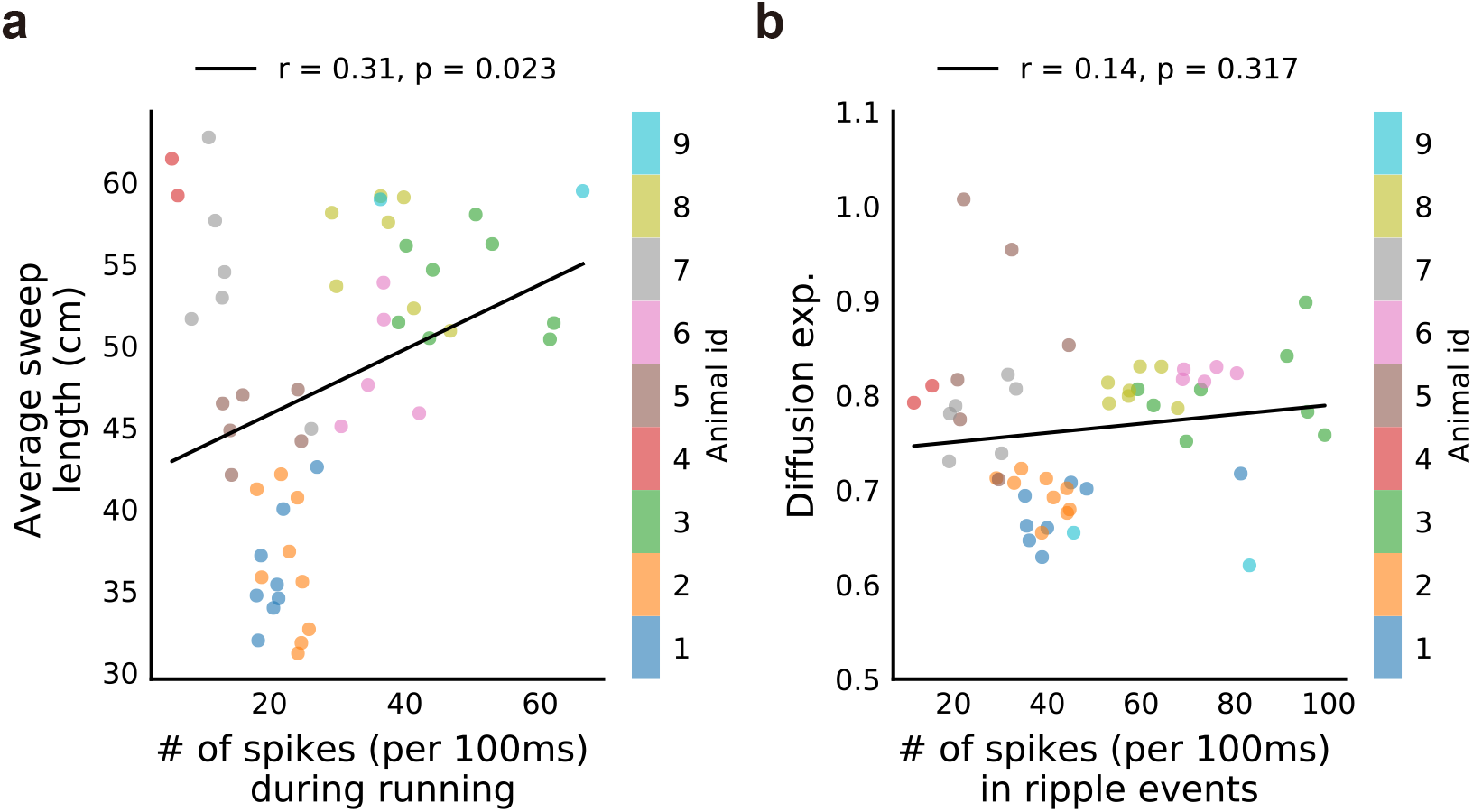
Correlation with the spike numbers participated in decoding. **(a)**, correlation between the theta sweep length and the number of spikes (per 100 ms) participated in online decoding (*r* = .31, *p* = 0.023, Pearson correlation). **(b)**, correlation between the diffusion exponent and the number of spikes (per 100 ms) participated in offline decoding (*r* = 0.14, *p* = 0.317, Pearson correlation).

**Figure S9:**
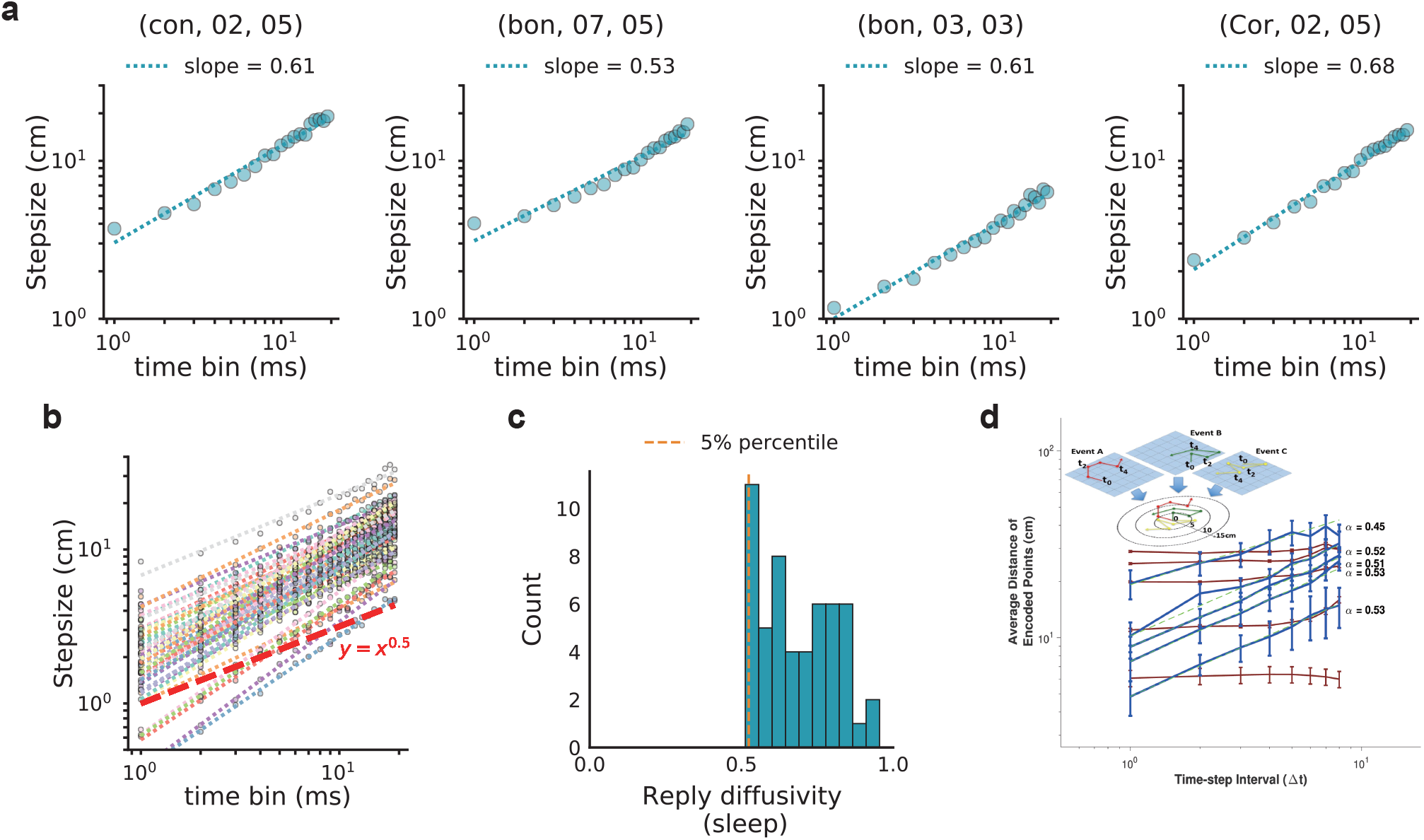
Diffusivity values during sleep replay. **(a)**, the log-log relationship between step size and time bin from four randomly selected sleep sessions. **(b)**, log-log plots from all valid sleep sessions. Each line represents one session (53 valid sessions in total). The red dashed line marks a typical Brownian-diffusion process with a slope of 0.5. **(c)**, histogram of diffusion exponents (one per sleep session) for sleep replay across all sessions and animals. The orange dashed line indicates the 5th percentile of the measured diffusivity values. Note that sleep replay is significantly larger than 0.5, diverging from the Brownian-diffusive dynamics. **(d)**, (adapted from Steele and Morris [1999] with permission) log-log plot of the time interval and the step size of replay trajectories for different mean reactivation speed groups. Note that the sleep replay reported in Steele and Morris [1999] follows the Brownian-diffusion dynamics with slope around 0.5.

